# Linguistic modulation of the neural encoding of phonemes

**DOI:** 10.1101/2021.07.05.451175

**Authors:** Seung-Goo Kim, Federico De Martino, Tobias Overath

**Author notes:** **Corresponding authors:** (S.-G. K.), (T. O.).

## Abstract

Speech comprehension entails the neural mapping of the acoustic speech signal onto learned linguistic units. This acousto-linguistic transformation is bi-directional, whereby higher-level linguistic processes (e.g., semantics) modulate the acoustic analysis of individual linguistic units. Here, we investigated the cortical topography and linguistic modulation of the most fundamental linguistic unit, the phoneme. We presented natural speech and ‘phoneme quilts’ (pseudo-randomly shuffled phonemes) in either a familiar (English) or unfamiliar (Korean) language to native English speakers while recording fMRI. This allowed us to dissociate the contribution of acoustic vs. linguistic processes towards phoneme analysis. We show that (1) the acoustic analysis of phonemes is modulated by linguistic analysis and (2) that for this modulation both of acoustic and phonetic information need to be incorporated. These results suggest that the linguistic modulation of cortical sensitivity to phoneme classes minimizes prediction error during natural speech perception, thereby aiding speech comprehension in challenging listening situations.

## 1 Introduction

Speech comprehension relies on the neural mapping of the acoustic speech signal onto linguistic categories (Hickok and Poeppel, 2007; Kleinschmidt and Jaeger, 2015; Poeppel et al., 2008). As such, the acoustic speech waveform that reaches our ears is converted into a neural code in the inner ear, which is then processed along the ascending auditory system and subsequently matched to learned linguistic categories (Friederici, 2011; Hickok and Poeppel, 2007). However, while this acousto-linguistic transformation is the basis for successful speech comprehension, many aspects of it still remain unknown. Here, we asked (1) whether the acousto-linguistic transformation is malleable to top-down linguistic information and (2) whether we can dissociate the contributions of acoustic and linguistic processing towards this transformation.

The phoneme is the smallest perceptual unit capable of determining the meaning of a word (e.g., the words *pin* and *chin* differ only with respect to their initial phonemes) (Stevens, 2000). Of the upward of 100 phonemes in use world-wide, approximately 44 phonemes make up the English language and these are categorized primarily based on articulatory features into four main classes: vowels, nasals and sonorants, plosives, fricatives and affricates (Ladefoged, 2001; Ladefoged and Johnstone, 2015). Each phoneme class has characteristic acoustic features; for example, while vowel sounds display a sustained period of harmonicity, plosives are characterized by a brief period of silence followed by a short broadband noise burst. Individual phonemes and the phoneme classes to which they belong have distinct temporal neural correlates: each phoneme class has a unique time-locked neural response characteristic, or phoneme-related potential (PRP; Khalighinejad et al. (2017); (Overath and Lee, 2017)). The phoneme-class-specific PRPs likely reflect the neural analysis of their acoustic characteristics (e.g., timing of energy onset, harmonicity, etc.) in functionally separate parts of auditory cortex.

In natural speech, phonemes do not occur in isolation, but instead form sequences to create syllables and words. The order in which phonemes can occur is governed by phonotactics, and is unique to each language (Chomsky and Halle, 1965). Apart from learning to recognize the language-specific phonemes themselves (Cheour et al., 1998), phonotactics is one of the first sets of rules infants need to learn during language acquisition (Friederici and Wessels, 1993; Jusczyk et al., 1994; Mattys and Jusczyk, 2001). This may be achieved via learning the likelihood of phoneme transitions: for example, in English certain phoneme transition probabilities are statistically unlikely (or even non-existent, e.g., /dla/) while others are statistically more likely (e.g., /gla/). A similar principle is thought to be employed for syllable transitions, where statistically improbable syllable transitions can indicate between-word boundaries (Saffran et al., 1996).

Thus, while the initial analysis of phonemes is based on their acoustic features (Khalighinejad et al., 2017; Mesgarani et al., 2014; Overath and Lee, 2017; Yi et al., 2019), subsequent processing stages are likely more linguistic in nature, such as those identifying language-specific phonemes or phonotactics, or even higher-level processes underlying the analysis of syntax, semantics, or lexical access (Friederici et al., 1993; Kocagoncu et al., 2017; Kutas and Hillyard, 1983). While decades of research have resulted in detailed speech/language models (Friederici, 2011; Hickok and Poeppel, 2007; Rauschecker and Scott, 2009), a clear demarcation between acoustic and linguistic analyses that contribute towards speech comprehension has largely remained elusive. One reason for this is that, in everyday listening situations, acoustic and linguistic analyses are difficult to separate and likely interact, e.g., via top-down modulation of acoustic feature analysis by linguistic processes (Anderson et al., 2003; Davis and Johnsrude, 2007; Díaz et al., 2008). In addition, previous studies that investigated phoneme processing in naturalistic contexts (Daube et al., 2019; Gwilliams et al., 2022; Khalighinejad et al., 2017; Mesgarani et al., 2014), did so only in a familiar language: this approach is unable to dissociate the initial acoustic processes from the obligatory nature of linguistic processes that become engaged in a native, familiar language.

In contrast, Overath and Lee (2017) were recently able to dissociate the acoustic and linguistic processes underlying phoneme analysis by comparing PRPs in familiar vs. foreign languages. They used a variant of a novel sound quilting algorithm (Overath et al., 2015) to create speech-based quilts in which linguistic units (phoneme, syllable, word) were pseudo-randomly ‘stitched together’, or quilted to form a new stimulus. This paradigm allowed the comparison of an acoustic stimulus manipulation (speech-based quilting) in a familiar vs. foreign language: if the processing of phonemes is affected by the acoustic manipulation (increasing linguistic unit size of speech quilts) in a familiar language only, then this would suggest that linguistic analysis in the familiar language influenced the acoustic analysis of phonemes. Put differently, if no phonemic repertoire or phonotactic rules are available to a listener (as is the case in a foreign language), the encoding of phonemes themselves should be independent of their ordering (phonotactics) or linguistic unit size in which they appear. Using EEG to investigate the PRP for different phoneme classes (Khalighinejad et al., 2017), Overath and Lee (2017) found that vowels in particular are amenable to such top-down linguistic modulation. However, the limited spatial resolution of EEG did not allow inferences as to where in the auditory cortex (or beyond) such top-down modulation might originate, or act upon.

Recent advances in fMRI time-series analysis have demonstrated that the neural activity to natural speech stimuli can be predicted from fast-paced acoustic (e.g., envelope, spectrum), phonological, and semantic features via linearized encoding modeling (De Heer et al., 2017; Huth et al., 2016). Inspired by this approach, the current study employed linearized encoding modeling of fMRI data in human cortex in an effort to reveal the separate encoding of acoustic and linguistic features of speech. Specifically, we used speech-based quilting (original speech vs. phoneme quilts) in familiar (English) vs. foreign (Korean) languages to dissociate the neural correlates of the acoustic and linguistic processes that contribute to the analysis of a fundamental linguistic unit, the phoneme. We show, for the first time, (1) that the acoustic analysis of phonemes is modulated by linguistic processes, and (2) that the interaction cannot be explained by solely acoustic or phonetic information.

## 2 Results

Ten native English speakers without any knowledge of Korean listened to speech stimuli in four conditions (original speech or phoneme quilts, in English or Korean) during three sessions of fMRI scanning. Condition-specific linearized encoding models were trained to predict the fMRI time-series using two “Acoustic” predictors (the broadband envelope and its first-order derivative with positive half-wave rectification) and four “Phonetic” predictors (the durations of each of the four main phoneme classes; i.e., vowels, nasals and approximants, plosives, fricatives and affricatives). Multi-penalty ridge regression models were optimized using a Bayesian optimizer for each predictor (i.e., 6 regularization hyperparameters per model). The prediction accuracy of full models with all predictors was calculated using Pearson’s correlation (*r*). The unique contribution of a certain predictor group (or a feature subspace; e.g., Daube et al. (2019)) was calculated using partial correlation (*ρ*) by regressing out the other predictor group (see **Figure 1** for an overview of the analysis).

**Figure 1.**
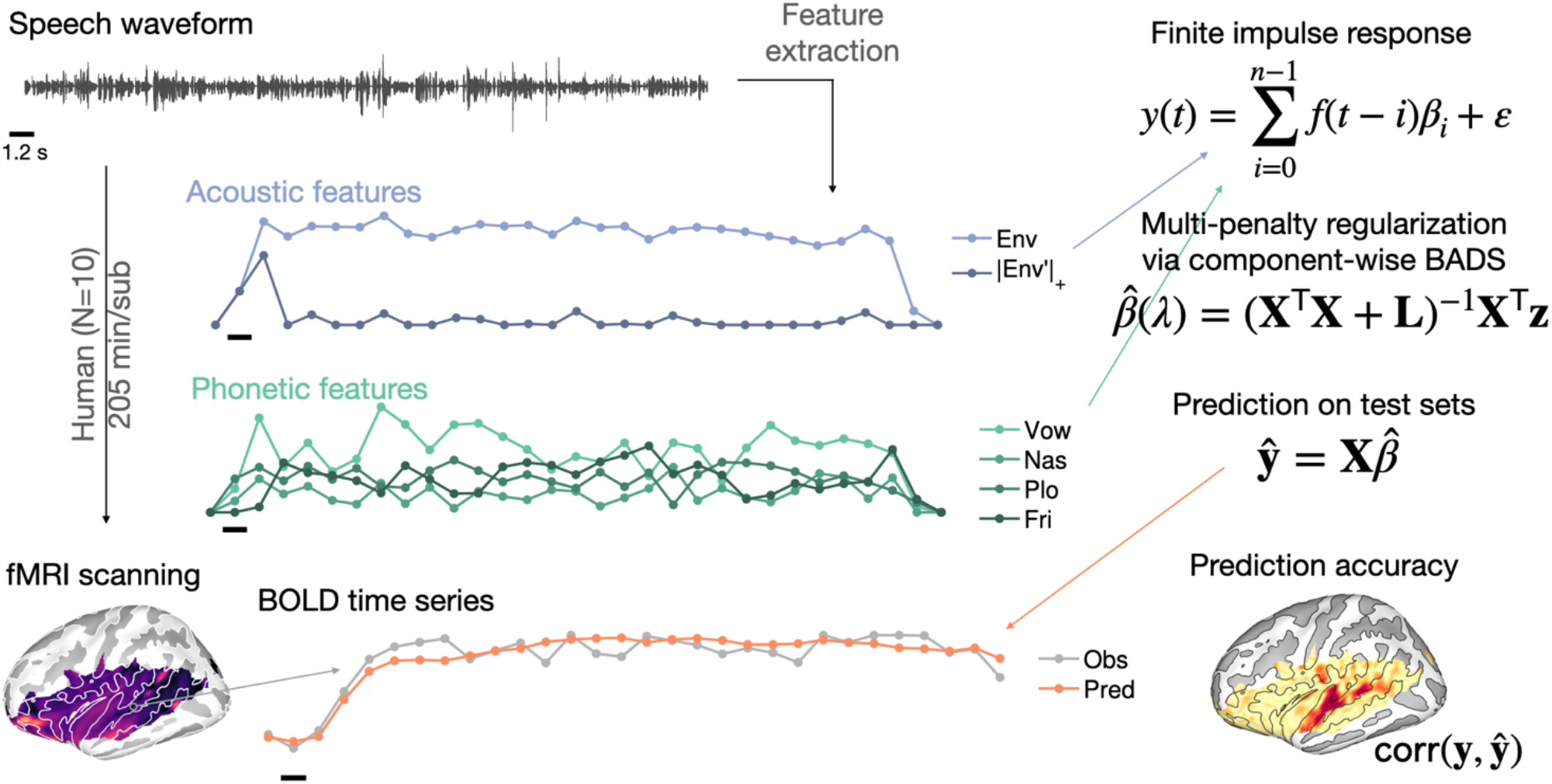
Linearized encoding analysis overview. Functional MRI data was acquired from 10 human participants while listening to unmanipulated or phoneme-scrambled speech stimuli in either English or Korean. From the speech waveform, acoustic features (cochleogram envelope and its first-order derivative with positive half-wave rectification) and phonetic features (the duration of four phoneme classes) were extracted and down-sampled at the fMRI sampling rate (1/1.2 Hz). Scale bars represent an fMRI sampling period (1.2 sec). After preprocessing, the surface-mapped BOLD time series y(t) was predicted using regularized finite impulse response modeling. Multi-penalty regularization was optimized in a principal component space using the Bayesian adaptive direct search (BADS) optimizer. The cross-validated prediction accuracy was measured by Pearson correlation between observed and predicted BOLD time series that was back-projected onto the vertex space (see Section 4.6 for details).

### 2.1 Linguistic processing interacts with acoustic processing in the left superior temporal sulcus

We first investigated whether the encoding models could replicate our previous findings of mean BOLD activity (Overath et al., 2015; Overath and Paik, 2021). **Figure 2** displays the differences in Pearson correlation of full models between conditions (see **Supplementary Figure S1** for a rendering on uninflated surfaces). The original speech stimuli evoked BOLD time-series that are better explained by the encoding models than the quilted stimuli in the superior temporal sulci (STSs) and the anterior superior temporal gyri (aSTGs), bilaterally (**Figure 2a**; max *t*[9] = 11.22, min cluster-*P* < 0.001, max cluster size = 1,078 vertices, max Δ*r* = 0.2302). The native language (English) as compared to the foreign language (Korean) showed similar but larger clusters over the lateral convexity of the STG (i.e., Te3; Morosan et al. (2005)), extending to the planum temporale in the left hemisphere (**Figure 2b**; max *t*[9] = 10.12, min cluster-*P* < 0.001, max cluster size = 1,697 vertices, max Δ*r* = 0.1913). An interaction in the expected direction (i.e., a greater difference for [Original > Quilts] in English than in Korean) was found in the superior portion of the left STS (i.e., Te4; **Figure 2c**; max *t*[9] = 7.98, min cluster-*P* = 0.002, max cluster size = 484 vertices, max Δ*r* = 0.1147), suggesting that the change of neural encoding as a function of the acoustic context was modulated by linguistic knowledge in this area.

**Figure 2.**
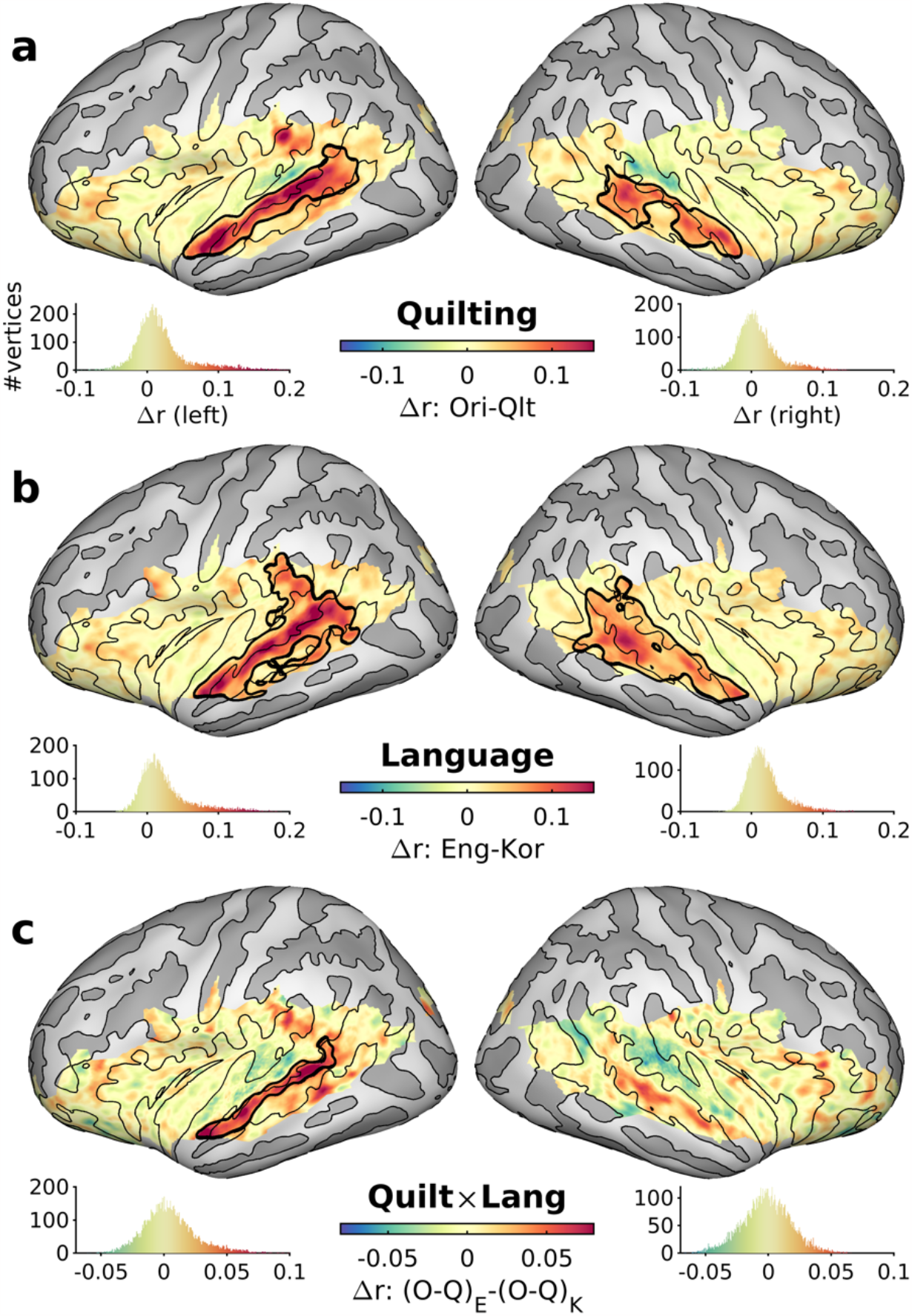
Comparisons of prediction accuracies between conditions: (a) main effect of Quilting, Original > Quilts; (b) main effect of Language, English > Korean; (c) interaction between Quilting and Language, [English-Original > English-Quilts] > [Korean-Original > Korean-Quilts]. Thick black contours mark significant clusters (cluster-P ≤ 0.005). Curvatures of the cortical surface are displayed in brighter (convex) and darker (concave) grays with thin black isocontours at the curvature of zero. Colored histograms of the *r* differences are displayed below each hemisphere. See Supplementary Figure S2 for comparisons at the subject-level.

**Figure 3.**
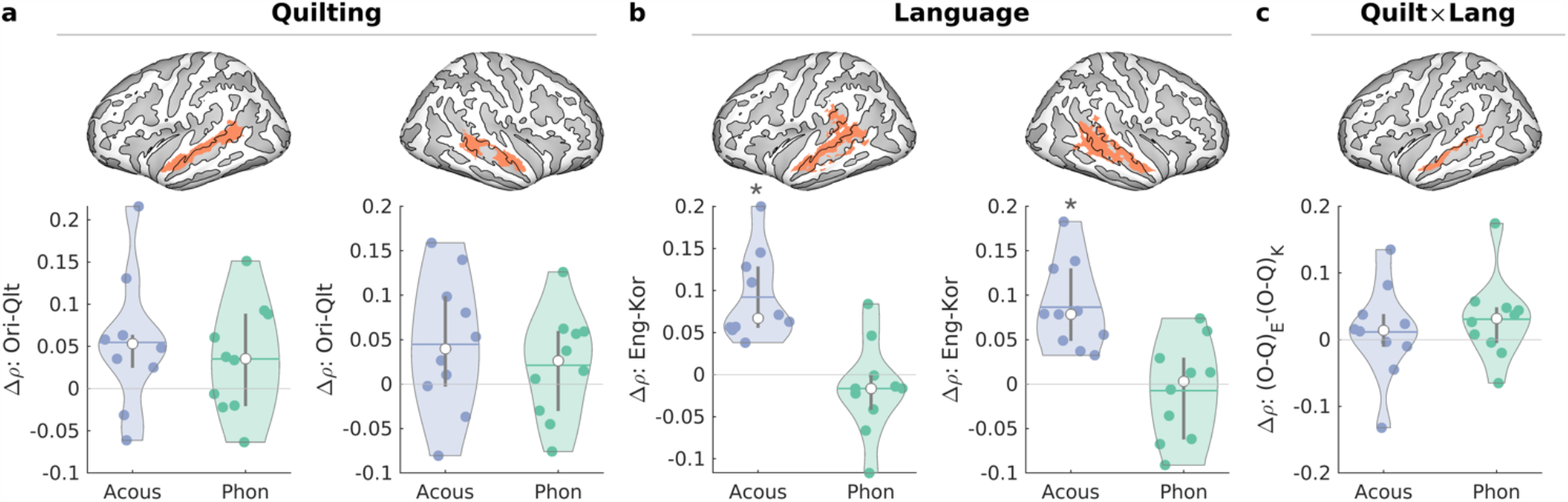
ROI-based partial correlation differences for contrasts (a) main effect of Quilting (Original > Quilts), (b) main effect of Language (English > Korean), and (c) interaction between Quilting and Language ([English-Original > English-Quilts] > [Korean-Original > Korean-Quilts]). Clusters are highlighted in orange. ROI-averaged partial correlations are shown in violin plots for the Acoustic (blue) and Phonetic (green) predictor groups; open circles mark medians, horizontal lines in colors mark means, and vertical gray thick lines mark the first and third quartiles. *: Bonferroni-corrected *P* < 0.005.

### 2.2 Neither acoustic nor phonetic predictors can exclusively explain the interaction between acoustic and linguistic processes

After establishing the interaction of acoustic and linguistic processes, we further examined whether the difference between original and quilted speech in the native language was driven by processes at the acoustic or phonetic level. An additional motivation for this analysis to separate acoustic and phonetic contributions was to account for the fact that the phoneme-based quilting procedure had slightly different effects on the envelope modulation spectrum in the two languages (**Supplementary Figure S3**). To this end, we calculated the partial correlation between the predicted and observed fMRI timeseries based on only one group of predictors (Acoustic or Phonetic), while regressing out the predicted timeseries based on the other group of predictors. The partial correlation thus quantifies the prediction based on the unique information in one predictor group in relation to the information that is common in both predictor groups. Finding an effect (of e.g., quilting) in the partial correlations would indicate that the linguistic modulation (i.e., the difference between original and quilted sounds in the native language as compared to the foreign language) is supported only by the unique characteristics of either Acoustic or Phonetic predictor groups that is orthogonal to common information of both predictor groups.

We tested whether the main effects and interaction identified in **Figure 2** can be explained by unique contributions of either of the predictor groups. For this, we averaged the partial correlation values in vertices within each cluster identified in **Figure 2**. This ROI-based comparison reveals that only for the main effect of Language, the partial correlation of the Acoustic predictors was significantly positive (max *t*[9] = 5.60, min *P* < 0.001, max diff *ρ* = 0.0921). That is, the residual STG/STS activity was better explained by the Acoustic predictors, but not by Phonetic predictors, for the native language (English) compared to the foreign language (Korean). For the interaction, neither of the predictor groups showed significant positive partial correlation differences (min *P* = 0.142), suggesting that the interaction shown in the full models in **Figure 2** cannot be explained by unique information of either of the two predictor groups, but is instead due to common information to both of them (see whole-cortex analyses in **Supplementary Figure S4**).

## 3 Discussion

The phoneme is the fundamental linguistic unit that determines the meaning of words. We show that the four main phoneme classes, as well as broadband envelopes, are encoded in fMRI data acquired while listening to continuous speech signals. The acoustic processes underlying this phoneme analysis are modulated by linguistic analysis, whereby the acoustic manipulation (original speech vs. phoneme quilts) affected speech encoding more in a familiar language than in a foreign language. The results also revealed that this modulation cannot be explained uniquely by either acoustic or phonetic predictors.

### 3.1 Linguistic modulation of acoustic analysis of phonemes

Our primary aim was to dissociate acoustic from linguistic processes, which would enable us to determine their interaction, i.e., whether linguistic processes modulate the acoustic analysis of phonemes. To this end, we found that the acoustic manipulation (phoneme quilts vs. natural speech) had a larger effect on phoneme processing in a familiar language (English) than in a foreign language (Korean). Since the acoustic manipulation was the same for both languages, this suggests that the greater difference between acoustic contexts was due to linguistic processes becoming engaged in a familiar language (however, see also Section 3.2 for further partitioning by predictor groups). Linguistic processes such as phonotactic, syntactic, as well as semantic analyses might therefore modulate the acoustic processing of phonemes, e.g., via hierarchical predictive coding or minimizing prediction errors through top-down modulation (Friston and Kiebel, 2009; Rao and Ballard, 1999). To our knowledge, this is the first demonstration of such linguistic modulation of a fundamental linguistic unit using fMRI. However, these results align well with Overath and Lee (2017), who found similar evidence for top-down linguistic modulation of phonemic analysis using a different recording modality (EEG).

Perhaps the best-known example of the modulatory influence of linguistic information is that of phonemic restoration (Samuel, 1981; Warren, 1970). In phonemic restoration, a phoneme is still subjectively ‘perceived’ even if it is masked or replaced completely by noise. This is often interpreted as an advantageous adaptation to speech perception in noisy environments, where it is common for interrupting or masking sounds to last only for a few tens or hundreds of milliseconds (i.e., on a temporal scale that is commensurate with that of phonemes). The top-down predictive nature of this phenomenon is further highlighted by the fact that, if the acoustic information is ambiguous, a ‘best guess’ phoneme is perceived (Leonard et al., 2016; Samuel, 1987). In fact, there is a wealth of evidence for such restorative processes in speech perception, for example from studies using noise-vocoded stimuli (Giraud et al., 2004; Narain et al., 2003; Obleser et al., 2008; Scott et al., 2000; Shannon et al., 1995; Wild et al., 2012) or other methods to distort the speech signal (Davis et al., 2011; Eckert et al., 2016), while the most common explanation for restorative effects refers to top-down, predictive (Friston and Kiebel, 2009) linguistic processes.

The locus of phonemic restoration, i.e. the region in which linguistic modulation is strongest, was recently shown to be situated in bilateral STG, likely due to receiving modulatory signals from left IFG (Leonard et al., 2016). This aligns remarkably well with the current study, where we found the strongest effect of linguistic modulation also along STG, albeit with a left-hemispheric dominance. The STG is a reasonable locus for such linguistic modulation, since it represents an intermediary processing stage in the language network that receives bottom-up information from primary auditory cortex and PT, as well as top-down information from higher-order auditory and frontal regions (Friederici, 2009, 2011; Hickok and Poeppel, 2007; Rauschecker and Scott, 2009). For example, the analysis of spectral shape (a necessary computation to differentiate between the formant structures of different vowels) relies on bottom-up changes in effective connectivity between HG to PT, as well as PT to STG/STS regions (Kumar et al., 2007; Warren et al., 2005). In contrast, top-down signals from frontal cortex (e.g., left IFG) have been shown to modulate speech processing in auditory cortex (Cope et al., 2017; Overath and Paik, 2021; Park et al., 2015; Sohoglu et al., 2012).

In the domain of electrophysiological measurements of speech perception, there is currently disagreement as to the extent that neural indices (such as speech-envelope entrainment, or phoneme encoding) can be interpreted as markers of linguistic processes that are necessary for speech comprehension (Di Liberto et al., 2015; Ding and Simon, 2013; Luo and Poeppel, 2007; Vanthornhout et al., 2018), or whether a more parsimonious explanation of these indices is that they reflect the analysis of characteristic acoustic properties of the speech signal (Baltzell et al., 2017; Daube et al., 2019; Howard and Poeppel, 2010; Millman et al., 2015; Verschueren et al., 2021). Our study is able to shed new light on this controversy by directly comparing the encoding of acoustic properties of phonemes in either a familiar language or in a foreign language, in which no higher-level linguistic analysis takes place.

We should note that the current study did not measure linguistic processes explicitly. For example, participants did not perform a linguistic task (e.g., speech comprehension), but were simply asked to detect a change in speaker, a task that is largely orthogonal to linguistic processing (see also Overath and Paik (2021) for a similar task). Therefore, we interpret the linguistic modulation of phoneme class analysis as obligatory linguistic processes that become engaged as soon as familiar linguistic templates (e.g., phonotactics, syntax, lexicon, semantics) are detected in the signal. Future studies will need to determine whether, and to what extent, these obligatory linguistic processes for phoneme analysis are malleable to various tasks that engage specific linguistic processes. For example, the neural processing of acoustic features in speech sounds has been shown to be enhanced or sharpened if they are task-relevant, attended to vs. ignored, or primed (Holdgraf et al., 2016; Leonard et al., 2016; Mesgarani and Chang, 2012; Rutten et al., 2019), and similar processes might become engaged for phoneme class encoding.

### 3.2 Contributions of acoustic and phonetic features

Another major advantage of the current study is the neural encoding analysis to delineate unique contributions of the overlapping information embedded in speech stimuli. The partial correlation analyses revealed a unique contribution of the acoustic features to the stronger neural encoding for the native language as compared to a foreign language (i.e., the main effect of language). Speech envelope (in particular onsets) encoding in the lateral STG has been shown from ECoG data with a native language (Oganian and Chang, 2019). As shown in the whole-cortex analyses (**Supplementary Figure S4**), this effect was strongest in the bilateral early auditory areas (HG/PT). This is in line with a recent finding that showed stronger speech envelope encoding in a native speaker group compared to a non-native speaker group with a low proficiency (Liberto et al., 2021).

However, no unique contribution of the acoustic or phonetic features was found for either the main effect of quilting or the interaction of language and quilting. This suggests that the stronger encoding of all features for the original than quilted speech samples (**Figure 2b)** and their interaction with language (i.e., more so for English than for Korean stimuli; **Figure 2c**) are driven by the shared information between the acoustic and phonetic predictors. A moderate multicollinearity between some predictors was indeed detected (see **Section 4.6.5**), presumably due to the similarity between the speech envelope and vowel predictors. Note, though, that each predictor was individually optimized using Bayesian optimizer to avoid suboptimal regularization for individual predictors; that is, it is likely that all individual predictors were optimally regularized in the present analyses. Taken together, the current fMRI data suggest that the shared information between envelope and vowel predictors supported the modulation of the encoding strength in the acoustic and linguistic contexts.

### 3.3 Encoding of envelope and phoneme classes in the BOLD time series

One of our preliminary aims was to confirm that rapid acoustic and phonetic features can be shown to be encoded in a hemodynamic response that is approximately two orders of magnitude slower (tens of milliseconds vs. seconds). Encoding of these features had previously been demonstrated using electro-/magneto-physiological methods, which afford commensurate millisecond temporal resolution (Di Liberto et al., 2015; Gwilliams et al., 2022; Khalighinejad et al., 2017; Yi et al., 2019). Nevertheless, the novel use of linearized ridge-regression modeling of fMRI BOLD signal time series was recently employed to successfully (and separably) reveal the encoding of acoustic and phonetic features in a familiar language: De Heer et al. (2017) collected fMRI data while presenting continuous, natural speech, and were able to reveal that the acoustic speech envelope predicted the BOLD time series best in HG, whereas articulatory phonetic features were predicted most accurately in higher-level auditory cortex such as STG. The current study is in broad agreement with these findings: while both the acoustic and phonetic features were encoded over the language network, the acoustic features showed greater encoding in the supratemporal regions (HG/PT).

More broadly, our study confirms that neural responses to rapid speech features, which are temporally integrated over several hundreds of milliseconds in the BOLD time series, can be revealed using linearized encoding modeling. Such models take advantage of the spatially separated functional organization of auditory cortex, for example with respect to prominent acoustic features such as frequency, spectro-temporal modulations, or spectral bandwidth (Baumann et al., 2015; Moerel et al., 2018; Rauschecker and Tian, 2004; Saenz and Langers, 2014; Santoro et al., 2014). This should encourage the future use of more naturalistic stimulus paradigms that allow the investigation of the complex dynamics of linguistic processes (Hamilton and Huth, 2020), as well as other higher-order processes such as music perception (Kim, 2022).

### 3.4 Modulation of acoustic and linguistic contexts

The analyses of the two factors Quilting and Language were motivated by previous studies that investigated the processing of temporal speech structure using segment-based speech quilting. In particular, these studies showed sensitivity in STS to temporal speech structure in either only a foreign language (Overath et al., 2015), or both native and foreign languages (Overath and Paik, 2021), which is comparable to a main effect of Quilting here. In addition, activity in left IFG revealed an interaction between Quilting and Language and increased as a function of temporal speech structure only in the familiar language (Overath and Paik, 2021). In the current study, Quilting and Language both had greater prediction accuracies in left STS, while their interaction in the same area was due to larger prediction accuracy differences for the Original vs. Quilts contrast in English vs. Korean.

For successful speech comprehension, the temporal dynamics of speech necessitate analyses at multiple scales that are commensurate with the average durations of phonemes, syllables, words, sentences, etc. This temporal hierarchy is thought to be reflected in a cortical processing hierarchy in which the neuronal temporal window of integration (Theunissen and Miller, 1995) increases from primary auditory cortex via non-primary auditory cortex to frontal cortex (e.g., (Lerner et al., 2011; Norman-Haignere et al., 2020); though see Blank and Fedorenko (2020) for a recent counterargument against such a hierarchy). The current results of greater prediction accuracy in STS as a function of Quilting largely support this view. A novel finding is the left-hemispheric lateralization. However, it is possible that this was driven by the interaction between Quilting and Language.

It is important to note that the segment-based quilting in previous studies disrupted the speech signal to a larger degree than the speech-based quilting employed here. The shortest segment length (30 ms) used in the previous studies, together with their placement irrespective of linguistic units, likely resulted in no phonemes being left fully intact in the resulting speech quilt. In contrast, the current speech-based quilting procedure preserved the phonemes (though likely still disrupted co-articulation cues).

### 3.5 Future directions

The current study makes a number of predictions for future studies investigating the acousto-linguistic transformation of speech. We show evidence for linguistic modulation of a fundamental linguistic unit, the phoneme, in native English speakers when listening to English speech, but not when listening to a foreign language for which participants had no linguistic repertoire. Therefore, while it is unlikely that the current results are specific to English phonemes, future studies should confirm this interaction, for example in native Korean participants who have no knowledge of English. Similarly, people who are perfectly bilingual in English and Korean should show evidence for linguistic modulation in both languages as a function of quilting, while those for whom both languages are foreign should not.

In addition, the fact that the linguistic modulation of the acoustic speech signal operates at an intermediate stage of linguistic analysis likely reflects its significance: if linguistic modulation starts at the level of phonemes, its ability to impact a later word processing stage is conceivably greater than if linguistic modulation only started at the word processing stage. Given the highly predictive nature of speech processing (see **Section 3.1** above), such modulation might be particularly helpful in situations in which the speech signal is compromised (e.g., in noisy conditions such as in a restaurant or bar). People with hearing loss (e.g., presbycusis) are a clinical population that is known to struggle in such situations, even with the help of hearing aids (Moore, 1996; Shinn-Cunningham and Best, 2008). It is therefore possible that (at least) one reason for their exacerbated speech comprehension difficulties in noisy situations is that the linguistic modulation of phonemes has deteriorated, thereby reducing the effectiveness of predictive speech processes. A similar argument might be made for people suffering from ‘hidden hearing loss’: i.e., hearing difficulties without detectable deficits in routine audiometry tests (Kujawa and Liberman, 2009; Ruggles et al., 2011). We predict that linguistic modulation of phoneme analysis is reduced in these populations (particularly in situations with background noise) and might thus serve as a clinical marker.

### 3.6 Conclusions

In conclusion, the current study demonstrates for the first time that individual phoneme classes derived from continuous speech signals are encoded in the BOLD signal time series. In particular, by using a design that dissociates acoustic from linguistic processes, we show that the acoustic processing of a fundamental linguistic unit, the phoneme, is modulated by linguistic analysis. The fact that this modulation operates at an intermediate stage likely enhances its ability to impact subsequent, higher-level processing stages, and as such might represent an important mechanism that facilitates speech comprehension in challenging listening situations.

## 4 Methods

### 4.1 Participants

Ten native English speakers without any knowledge or experience in Korean participated in the current study (mean age = 24.0 ± 2.2 years; 6 females). Eight participants volunteered in three sessions consisting of 8 runs each on separate days (intervals in days: mean = 8.5, standard deviation = 16.6, min = 1, max = 70) and two other participants in a single session each (6 runs and 8 runs, respectively), resulting in a total of 24 scanning sessions. This is on par with similar approaches that maximize intra-subject reliability over intra-subject variability in the data (Breedlove et al., 2020; Huth et al., 2016; Kay et al., 2008; Moerel et al., 2013; Naselaris et al., 2015; Norman-Haignere et al., 2015; Santoro et al., 2017).

All participants were recruited via the Brain Imaging and Analysis Center (BIAC) at Duke University Medical Center, NC, USA after safety screening for MRI (e.g., free of metal implants and claustrophobia). All reported to have normal hearing and no history or presence of neurological or psychiatric disorders. Informed written consent was obtained from all participants prior to the study in compliance with the protocols approved by the Duke University Health System Institutional Review Board.

### 4.2 Stimuli

Speech stimuli were created from recordings (44,100 Hz sampling rate, 16-bit precision) of four female bilingual (Korean and English) speakers reading textbooks in either language as in previous studies (Overath and Lee, 2017; Overath and Paik, 2021). Native English and Korean speakers judged the recordings as coming from native English and Korean speakers, respectively. Korean was chosen because of its dissimilarity to English: it shares no etymological roots with English and has different syntactic and phonetic structures (Sohn, 2001).

We used a modified version of the quilting algorithm (Overath and Lee, 2017; Overath and Paik, 2021) where we pseudorandomized the order of phonemes (instead of set segment lengths). First, phonemes were extracted from the recordings and corresponding transcripts using the Penn Phonetic Lab Forced Aligner^1^ (Yuan and Liberman, 2008) for English speech and the Korean Phonetic Aligner^2^ (Yoon and Kang, 2013) for Korean speech. The phoneme segmentation output was a Praat TextGrid, which was then imported to MATLAB^3^ via the mPraat toolbox (Bořil and Skarnitzl, 2016). The alignment was manually validated by a native English and Korean speaker, respectively (Overath and Lee, 2017; Overath and Paik, 2021). The durations of phonemes in the recordings of natural speech in milliseconds were as follows (see **Supplementary Figure S5a** for histograms): min = 4.3, max = 396.2, mean = 72.8, median = 63.8, standard deviation = 41.7, skewness = 1.2 in English (*N* = 10,514); min = 8.9, max = 308.3, mean = 71.9, median = 63.7, standard deviation = 36.0, skewness = 1.3 in Korean (*N* = 10,894). The average durations were similar between languages (0.9 ms longer in English, *t*[21406] = 1.67, *P* = 0.094) while the distributions were slightly different for that English had more instances of short (e.g., < 20 ms) phonemes (Kolmogorov-Smirnov statistic = 0.1413, *P* = 10^−93^).

The phoneme segments were pseudorandomly rearranged to create novel phoneme quilts. For each stimulus, a random initial phoneme was chosen; subsequent phonemes were selected such that the acoustic change at the boundary was as close as possible to the acoustic change in the original source signal (using the L2-norm metric of an ERB-spaced cochleogram; see (Overath et al., 2015)). In addition, we applied the following exclusion criteria: (a) the phoneme duration needed to be at least 20 ms, (b) two identical phonemes could not occur next to each other, (c) for a given phoneme, its subsequent phoneme could not be the same as in the original source signal. We used the pitch-synchronous overlap-add (PSOLA) algorithm (Moulines and Charpentier, 1990) to further minimize abrupt changes in pitch at phoneme boundaries. Overall alterations due to the quilting algorithm were quantified by the Kullback-Leibler divergence (*D*_KL_) between L2-norm acoustic change distributions in the original source and the created phoneme quilt (median *D*_KL_ = 0.6873 bits for English, 0.6004 bits for Korean; Wilcoxon rank sum equal median test: *Z* = 0.5913, *P* = 0.5543). In the phoneme quilts, the durations of phonemes in milliseconds were as follows (**Supplementary Figure S5b**): min = 20.0, max = 351.0, mean = 72.3, median = 63.0, standard deviation = 39.4, skewness = 1.4 in English (*N* = 10,467); min = 20.0, max = 383.0, mean = 69.7, median = 60.0, standard deviation = 36.3, skewness = 1.5 in Korean (*N* = 11,213). There were slight differences between languages in average durations (2.6 ms longer in English, *t*[21678] = 5.08, *P* = 10^−7^) and distributions (KS-stat = 0.0657, *P* = 10^−21^), however, the mean difference of 2.6 ms is much shorter than the modeled cochlear integration time-window of 20 ms.

As for the temporal modulation of the speech stimuli (**Supplementary Figure S3**), the phoneme-based quilting decreased the temporal modulation energy at around 3–10 Hz in both languages. However, this reduction was greater in Korean than in English (max *F*[1,23] = 41.09; min FDR-*P* = 0.0005; significant [FDR-*P* < 0.05] frequency bins = 3.9, 5.0, 5.1, 5.7, 6.4, 6.5, 6.6 Hz, in total 7 frequency bins; max effect size = 5.11 dB). This motivated us to include acoustic predictors in the encoding analysis (see **Section 4.6.2**).

For both languages (English and Korean), the 33-s long stimuli in the two experimental conditions (Original and Phoneme Quilts) were created by concatenating six 5.5-s stimuli (24 unique exemplars per condition and language). Subsequent 5.5-s stimuli were either from the same or a different speaker (participants were asked to detect changes in the speaker, see **Section 4.3**). The overall sound intensity was normalized by equalizing the root-mean-square (RMS) signal intensity across stimuli. At the beginning and the end of the 33-s stimuli, 10-ms cosine ramps were applied to avoid abrupt intensity changes.

### 4.3 Experimental procedure

Functional MRI data were acquired while participants listened to the speech stimuli (either Original or Phoneme Quilts in either language) and performed a task to maintain attention to the stimuli. A 33-s trial consisted of six 5.5-s stimuli of multiple speakers in a given condition. Silent inter-stimulus intervals (ISIs) were uniformly varied between 5.6 s and 10.4 s (mean = 8 s). One run consisted of twelve 33-s trials, and one session consisted of eight 8.5-min runs (except for one participant, who only completed six runs). For one of the eight participants with three sessions, one run was prematurely terminated after 9 of the 12 trials due to technical difficulties (the intact 9 trials from the run were still used in the analysis). In total, fMRI data corresponding to ∼203 min/participant were obtained for the 8 participants with 3 full sessions (average of ∼174 min/participant for all 10 participants); this corresponds to ∼158 minutes of stimuli (excluding the ISI) per participant with 3 full sessions (average of ∼137 min/participant for all 10 participants).

The stimulus presentation timing was controlled via the Psychophysics Toolbox (v3.0.11^4^). Each run was triggered by the TTL signal from the MRI scanner mediated by a counter. Digital auditory signals at 44,100 Hz sampling rate and 16-bit precision from a Windows desktop were converted to analog signals by an external digital amplifier (Sony, Tokyo, Japan) and delivered to participants via MRI-compatible insert earphones (S14, Sensimetrics, MA, USA) at a comfortable listening level (∼75 dB SPL). Participants wore protective earmuffs on top of the earphones to further reduce acoustic noise from the MRI scanner.

The task was to indicate a change in speaker (i.e., a 5.5. s stimulus of one speaker followed by a different speaker) via a button press on an MRI-compatible four-button pad (average speaker changes per trial = 3.5, between 1 and 4). The performance was assessed via d-prime *d′* = Φ^−1^(Pr(*Y*|*s*)) – Φ^−1^(Pr(*Y*|*n*)) where Pr(*Y*|*s*) is the hit rate in “signal” trials and Pr(*Y*|*n*) is the false alarm rate in “noise” trials and Φ^−1^(·) is the inverse cumulative distribution function of the zero-mean, unit-variance Gaussian distribution (Macmillan and Kaplan, 1985). Responses were classified as a hit if they occurred within 3 s following a change in speaker (and otherwise classified as false alarm). In the case of multiple responses within one 5.5-s stimulus segment, only the first response was counted. For extreme values of hit/false alarm rates (i.e., 0 or 1), an adjustment (i.e., adding 0.5/n to zero or subtracting 0.5/n from one for n trials) was made to avoid infinite values of *d’* (Macmillan and Kaplan, 1985).

After each 33-s trial, participants received visual feedback about their performance (*D′* = *d’/max d′ where max d’* is a *d’*for a perfect performance, ranging between [-100%, 100%]) with a description (“POOR” for *D’* < 0, “FAIR” for 0 ≤ *D’* < 50%, “GOOD” for 50% ≤ *D’* < 100%, “PERFECT!” for *D’* = 100%) to encourage continued attention. While multiple button presses were discarded from computing d’, an alerting message was presented to the participants (“NO KEY PRESSED!” or “TOO MANY KEYS PRESSED!”) instead of the performance feedback when the button presses were too many (> 5) or none (2.5% of total 2,397 trials from 9 participants; participant 1 was excluded from the d-prime analysis due to a technical fault of the in-scanner response device). The average *D’* was 61.1% ± 38.4% points (d’ = 1.14 ± 0.72), without a significant difference between languages (repeated-measures ANOVA, 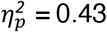, *F*[1,7] = 5.46, *P* = 0.21), but between original speech and phoneme quilts (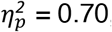, *F*[1,7] = 16.37, *P* = 0.02).

### 4.4 Image acquisition

All images were acquired using a GE MR 750 3.0 Tesla scanner (General Electric, Milwaukee, WI, USA) with a 32-channel head coil system at the Duke University Hospital, NC, USA. For blood-oxygen-level-dependent (BOLD) contrast, gradient-echo echo-planar imaging (GE-EPI) with a simultaneous multi-slice (SMS) acceleration factor of 3 (i.e., 3 slices acquired in parallel with aliasing of FOV/3 shifts) was used (in-plane pixel size = 2 × 2 mm^2^, slice thickness = 2 mm, slice gap = 0 mm, FOV = 256 mm, matrix size = 128 × 128, TE = 30 ms, flip angle = 73°, TR = 1200 ms, phase-encoding direction = posterior-to-anterior). A total of 39 slices were acquired for each volume (13 slices per band) in an interleaved ascending sequence. At the beginning of a run, the volume was centered on the supratemporal plane, covering from the inferior colliculus to the inferior frontal gyrus. To correct for magnetic inhomogeneity artifacts, an additional GE-EPI image of 3 volumes with a reversed phase encoding direction (anterior-to-posterior) was acquired after each run except for the first participant.

For T1-weighted contrast, a magnetization prepared rapid gradient echo (MP-RAGE) scan covering the whole-brain (in-plane pixel size = 1 × 1 mm^2^, slice thickness = 1 mm, slice gap = 0 mm, FOV = 256 mm, matrix size = 256 × 256, TE = 3.2 ms, flip angle = 8°, TR = 2264 ms, number of slices = 156) was acquired at the end of each session.

### 4.5 Image processing

#### 4.5.1 Anatomical images

T1-weighted images were segmented using SPM (SPM12; v7487^5^) to obtain tissue probability maps (*spm.spatial.preproc*), which were used for anatomical CompCor regressors (Behzadi et al., 2007). High-resolution cortical surfaces were fully automatically constructed using FreeSurfer (v6.0.0^6^) for surface-based analysis.

#### 4.5.2 Functional images

The displacement due to inhomogeneity in the B0 field (i.e., susceptibility artifacts) was corrected using *topup* in FSL (v5.0.11^7^) with the reversed phase-encoding images. The first 6 volumes (i.e., “dummy scans”) of each run were subsequently discarded from the analyses. Temporal and spatial realignments were achieved using SPM: the slices were first temporally aligned to the center of the TR using sinc-interpolation (*spm.temporal.st*), and then the volumes were spatially aligned to the mean volume using 4-th degree B-spline interpolation (*spm.spatial.realignunwarp*). Since we used a multiband sequence (i.e., 3 slices were acquired simultaneously), the acquisition time of each slice and reference time were provided (instead of slice order) for the slice-timing correction.

Anatomical CompCor regressors were extracted from realigned EPI volumes. On concatenated time series from voxels with > 99% probability for white matter and cerebrospinal fluid, principal component analysis (PCA) was applied to extract principal components. Six components with highest eigenvalues were used as “CompCor” regressors in the GLM denoising procedure (see **Section 4.5.3**).

Next, the EPI volumes were projected onto individual cortical surfaces (∼150,000 vertices per hemisphere) at the middle depth of cortices by averaging samples at the 40%, 50%, and 60% of cortical thickness to avoid aliasing (*mri_vol2surf* in FreeSurfer). Surface-mapped functional data were normalized to ‘fsaverage6’ surfaces (40,962 vertices per hemisphere) via spherical surface registration, and then smoothed with a 2-D Gaussian kernel with the full-width-at-half-maximum (FWHM) of 6 mm (i.e., 3 pixels in the EPI slices) via iterative nearest-neighbor averaging (*mri_surf2surf* in FreeSurfer).

#### 4.5.3 Surface-based GLM denoising

We applied a model-based denoising technique for task-based fMRI data (GLMdenoise v1.4^8^) to the surface-mapped data (Kay et al., 2013). The algorithm extracts ‘noise’ regressors from the data that would increase prediction accuracy in leave-one-run-out-cross-validation. This is achieved by first defining ‘noise pool’ voxels with negative R^2^ values for a given design matrix (i.e., voxels that are irrelevant to the task of interest), extracting principal components from the noise pool, and then determining an optimal number of components to remove as a minimal number where the improvement in cross-validation prediction decays. We used box-car functions to represent the four conditions in the design matrix. On average, 4.5 ± 2.1 noise regressors were regressed out. These improved reliability in estimation (mean over standard errors ratio of coefficients estimates across CV folds: median increase = 0.82; mean increase = 1.12) but only slightly increased predication accuracy (cross-validation R^2^: median increase = 0.25% points; mean increase = 0.56% points). In addition to the noise regressors, the 4-th order polynomial fits to slow drifts in BOLD time series, the six CompCor regressors, and the button-press regressors convoluted with a canonical HRF were regressed out from the residuals (i.e., prediction from the design matrix subtracted from the data).

### 4.6 Linearized encoding analysis

We predicted BOLD time series at each vertex in response to speech sounds using a linearized encoding model based on finite-impulse response (FIR) functions. Multiple lags were used to model the variable hemodynamic responses in different cortical areas (De Heer et al., 2017; Huth et al., 2016). In order to account for the collinearity of predictors representing acoustic and phonetic information, we used ridge regression to fit the model (i.e., FIR weights) and evaluated the prediction via cross-validation. The procedures are explained in detail in the following subsections.

#### 4.6.1 Vertex selection

For our interest in auditory and linguistic processing, we restricted our analysis to vertices in cortical regions that are previously known to be involved in speech processing so as to avoid unnecessary computations. Specifically, from the automatic parcellation based on the Desikan-Killiany cortical atlas (Desikan et al., 2006), the following 19 labels were included: ‘bankssts’, ‘caudalmiddlefrontal’, ‘inferiorparietal’, ‘inferiortemporal’, ‘lateralorbitofrontal’, ‘middletemporal’, ‘parsopercularis’, ‘parsorbitalis’, ‘parstriangularis’, ‘postcentral’, ‘precentral’, ‘rostralmiddlefrontal’, ‘superiorparietal’, ‘superiortemporal’, ‘supramarginal’, ‘frontalpole’, ‘temporalpole’, ‘transversetemporal’, ‘insula’. Selected regions are visualized in **Supplementary Figure S6**. Vertices with BOLD time series varied across participants due to the variability of head sizes, individual acquisition volumes at each session, and movements across runs during sessions. **Supplementary Figure S7** shows the overlap of selected vertices across participants. On average, 28,297 ± 3,748 vertices were selected per participant.

#### 4.6.2 Predictors

We included as predictors (i) the durations of phoneme classes (vowels, nasals and approximants, plosives, fricatives and affricatives; Vo, Na, Pl, Fr, respectively), and (ii) the speech envelope and its first-order derivative with positive half-wave rectification. For (i), the onset time and duration of each phoneme were determined and then grouped according to phoneme class (Ladefoged and Johnstone, 2015; Shin, 2015) (see **Supplementary Table S1**). Bigram transition probabilities between phoneme classes (**Supplementary Figure S8**) were effectively altered by the quilting algorithm (Hotelling’s *T*^2^ between Original and Phoneme-quilts = 1563, *P* < 10^−6^ for English; Hotelling’s *T*^2^ = 1258, *P* < 10^−6^ for Korean). The durations of phoneme classes were modelled as box-car functions at the audio sampling rate (44.1 kHz) and were then down-sampled to 1/TR (1/1.2 = 0.833 Hz) following anti-aliasing low-pass filtering. To align with the slice timing correction applied to the BOLD time series, the resampled time points were also at the center of the TR. For (ii), the speech envelope was computed from a cochleogram (30 filters from 20 to 10,000 Hz, equally spaced on an equivalent rectangular bandwidth [ERB] scale) by raising the Hilbert envelope of the resulting cochleogram to a power of 0.3 to simulate cochlear compression and summing energy across all 30 ERB channels (McDermott and Simoncelli, 2011; Overath et al., 2015). The speech envelope was then down-sampled as for the phoneme class durations. The rectified derivative was calculated following the down-sampling to reflect slow temporal modulations.

The down-sampled predictors showed moderate collinearity; the diagnostic metric of the collinearity, namely “condition index”, had an overall value of 27 (23–24 in English conditions and 41 in Korean conditions) given a suggested criterion (> 30) for a moderate multicollinearity (Belsley, 1991). The condition index is a square root of the maximum eigenvalue divided by the minimum eigenvalue of the design matrix, quantifying the upper bound of the collinearity of the design matrix. The observed multicollinearity was mainly due to the high dependency between the vowel and the envelope predictors; the proportions of explained variance by the corresponding eigenvector (i.e., variance decomposition proportion; VDP) were 0.87 and 0.99 for the vowels and the envelope, respectively. The collinearity patterns were similar across conditions (**Supplementary Figure S9**). The existence of multicollinearity motivated the use of a penalized regression.

#### 4.6.3 Regularized finite-impulse response modelling

A FIR model was used to predict the BOLD time series at each vertex. In this approach, we modelled the neural response as a convolution of the predictors and a linear FIR filter, which is a commonly used approach in receptive field mapping of neural populations (Ringach et al., 1997; Wu et al., 2006).

Consider a linear model for *t* time points and *p* predictors,

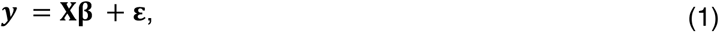

where ***y*** is a (*t* × *1*) data vector (i.e., BOLD time series at a certain voxel), **X** is a (*t* × *p*) design matrix (i.e., a FIR model), **β** is a (*t* × *p*) unknown coefficient vector, and **ε** is a noise vector from a zero-mean Gaussian distribution with a serial correlation ε∼𝒩(**0**, *σ*^*2*^ Ω) where Ω is a (*t* × *t*) unknown covariance matrix and *σ*^*2*^ is a scale factor. For the FIR modeling, the design matrix **X** consists of matrices of delayed features as:

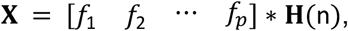

for *p* features and *n* delays as implemented in a convolutional kernel **H**(n), while ∗ denotes the convolution operation. A Toeplitz matrix can be constructed for delayed features between time point *t*_*1*_ and *t*_*2*_ with *n* delays for the *i*-th feature as:

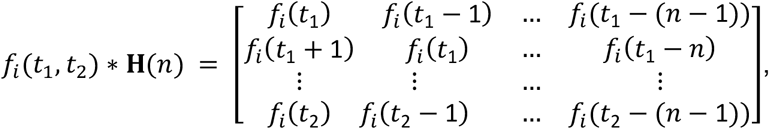

where *f*_*i*_(*t*) is the scalar value of the *i*-th predictor at time point *t*. In the current study, we delayed the predictors by 1, …, 10 TRs (1.2, …, 12 s). Once unknown coefficients (or weights) are estimated, an inner product 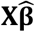 is effectively a convolution of the *i*-th feature and the estimated filter.

While it is standard to pre-whiten the data when modeling autocorrelated noise for a Generalized Least Squares (GLS) solution (Aitken, 1936), here we did not pre-whiten the model. This is because even with autocorrelated noise, an Ordinary Least Squares (OLS) solution is still an unbiased estimator (only its efficiency is suboptimal) and because our goal was to estimate (predict) responses, not to infer significance. In particular, for the current data, GLS often yielded worse cross-validation prediction than OLS. Therefore, we empirically determined not to pre-whiten the model.

As we detected a strong collinearity among the predictors, we applied L_2_-norm regularization to solve Eq. (1), which is known as a multi-penalty ridge solution (Hoerl and Kennard, 1970):

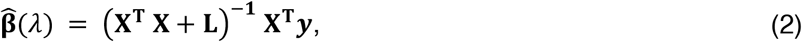

where 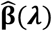 is a vector of penalized estimates and **L** is a regularization matrix as:

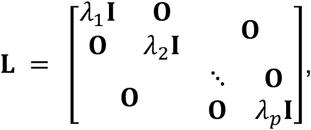

with λ_*i*_ is a scalar regularization hyperparameter for the *i*-th feature, **I**_*i*_ is the (*n* × *n*) identity matrix, and is a zero matrix in appropriate dimensions. The multi-penalty ridge has been recently re-introduced to the neuroscience community as “banded ridge” (Nunez-Elizalde et al., 2019).

#### 4.6.4 Component-wise optimization and voxel-wise evaluation

We used nested cross-validation (CV) to optimize and evaluate the models (Hastie et al., 2009). To avoid information leakage driven by the stimulus-evoked responses (Hasson et al., 2010; Kaufman et al., 2012), trials were partitioned in a way that there is no overlap of stimuli across partitions. The whole data was partitioned into two outer-CV folds (training set and test set) to evaluate model performance on unseen data; the outer-CV training set is further partitioned into two inner-CV folds (training set and validation set) to optimize hyperparameters on independent data (Varoquaux et al., 2017).

Optimizing multiple penalty terms can be a non-trivial task (van de Wiel et al., 2021). Grid search algorithm can be efficiently implemented using general linear model (GLM): i.e., a single inversion of the regularized design covariance matrix (**X**^T^**X** + **L**) can be used for all vertex-wise models for a particular search value. However, with the increasing number of penalty terms, the combinations of search values exponentially increase. Bayesian adaptive direct search (BADS) optimizer^9^ is known to be robust in optimizing high-dimensional parameters (Acerbi and Ma, 2017). But, as the BADS algorithm finds a unique optimization path in the parameter space, it runs through unique combinations of search values for each initialization. That is, separate inversions of differently regularized covariance matrices are required for each model; this process cannot be done simultaneously across all vertices (models) unlike the grid search using GLM.

To reduce the number of models to optimize, we exploited the spatial dependency of fMRI and data dimensionality (i.e., much fewer time-points than vertices). Using PCA on temporally concatenated data, we transformed the vertex time-series into the component space as:

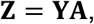

where **Z** is a (*T* × *k*) component time-series matrix, **Y** is a (*T* × *v*) vertex time-series matrix for *T* time-points (with all trials are temporally concatenated) and *v* vertices (28,297 ± 3,748 on average), **A** is a (*v* × *k*) demixing matrix for *k* components, which was determined for each subject to explain 99% of the total variance of the vertex time-series (1,486 ± 152 on average; 5.25% of the number of vertices). Instead of a vertex-wise model (Eq. 2), we fitted a component-wise model as:

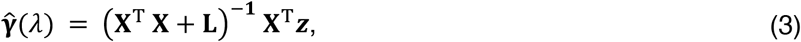

where **γ** is a coefficient vector for a component and ***z*** is a component time-series for a given component. The BADS optimizer searched the exponents of base 10 for λs with absolute bounds of [-15, 15] and plausible bounds of [-10, 10]. Over 10 random initializations (uniformly sampled within the plausible bounds), a vector of exponents that minimizes the validation error (sum of squares) was chosen for each component. The optimization process took 79–102 hours per outer-CV fold, depending on the number of sessions, on Intel Xeon Gold 6130 processors.

For evaluation, we predicted the component time-series for the outer-CV test set using the weights estimated from the outer-CV training set: 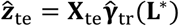 where subscripts ‘tr’ and ‘te’ indicate the outer-CV training set and test set, respectively, and **L**\* denotes the optimal regularization matrix. Then, the predicted component time-series was transformed back into the vertex space: 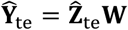 where **W** = **A**^**−1**^ is a (*k* × *v*) mixing matrix. Finally, for each vertex, the prediction accuracy was calculated by Pearson’s correlation (*r*) between the observed time-series and the predicted time-series: *r* = corr(**y**_te_, **ŷ**_te_).

To determine the uniquely explained variance by a particular set (subspace) of predictors (either the Acoustic or Phonetic subspace), vertex time-series were separately predicted based on each subspace as:

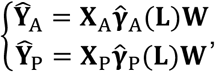

where **X** = [**X**_A_ **X**_P_] and **γ** = [**γ**_A_ **γ**_P_]. Note that the sum of separate predictions is equal to the full model prediction **Ŷ** = **Ŷ**_A_ **+ Ŷ**_P_. Then, partial correlation (*ρ*) was calculated as Pearson’s correlation between residuals after regressing out one prediction from the data and the other prediction as *ρ*_A_ = corr(**y, ŷ_A_**; **ŷ**_P_) and *ρ*_P_ = corr(**y, ŷ**_P_; **ŷ**_A_).

#### 4.6.5 Statistical inference

Statistical inference was computed via a non-parametric paired *t*-test using a cluster-based permutation test at group-level (Maris and Oostenveld, 2007). Specifically, *r* values of both models were calculated for each participant (N = 10), and then the difference between two models at each vertex was calculated. Next, the signs of differences across participants were flipped over all possible permutations (2^10^ = 1,024) to form a null distribution. One-tailed *P*-values were computed from the null distribution for our directed hypotheses (e.g., English > Korean, Original > Quilts). Note that the inference was computed at the group-level, not the subject-level. Vertex-wise multiple comparisons correction was applied via a cluster-based permutation test as implemented in *ft_statistics_montecarlo* in FieldTrip (v20180903)^10^ with a custom modification of *clusterstat* for a faster cluster identification through parallelization. In an earlier fMRI methodological study (Eklund et al., 2016), it was shown that a liberal cluster-forming threshold (CFT) in a cluster-level inference based on the random field theory resulted in a severely inflated family-wise error rate (FWER), whereas the permutation test showed a consistently proper control of the FWER regardless of the choice of a CFT. A recent study formally showed that a CFT in permutation tests does not affect the FWER, but only the sensitivity (Maris, 2019). Thus, in the current study, clusters were defined by an arbitrary threshold of the alpha-level of 0.05 (for vertex-wise *P*-values) to improve the sensitivity, and the cluster-wise *P*-values are thresholded at the alpha-level of 0.005 to control the FWER at 0.005.

### 4.7 Data availability

The data that support the findings of this study are available from the corresponding authors upon reasonable request.

### 4.8 Code availability

The computer code that was used for this study is available on the Open Science Framework repository^11^.

## 5 Acknowledgements

The authors would like to thank Frankie Pennington and Joon Hyun Paik for manually checking phoneme onsets and offsets for the forced alignment of English and Korean recordings, respectively. This work was supported by US National Institutes of Health grant R21DC016386 to T.O.

## 7 Supplementary materials

### 7.1 Supplementary figures

**Figure S1.**
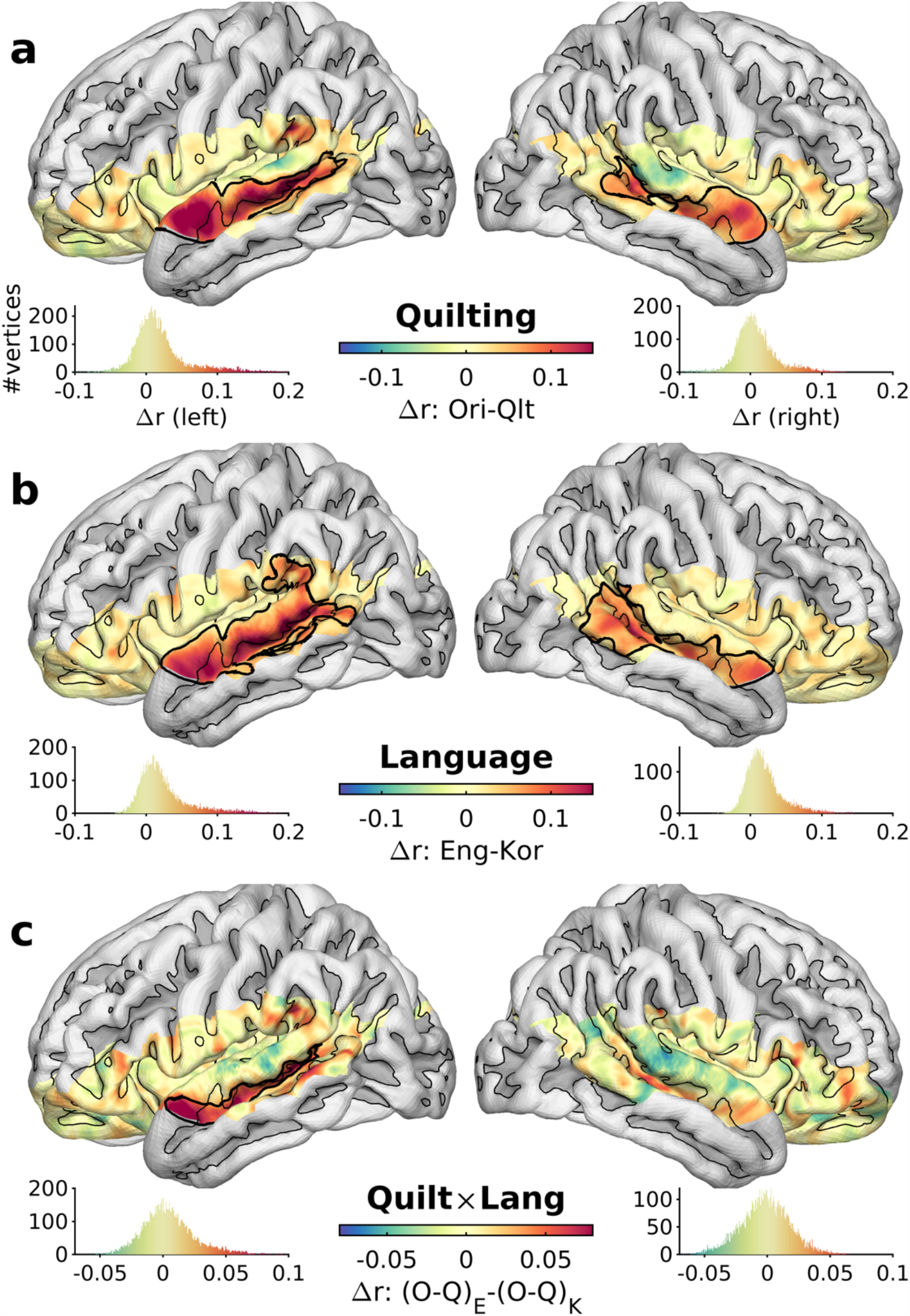
Pial surface rendering of Figure 1 for anatomical interpretation. The same schematics as Figure 1 are used.

**Figure S2.**
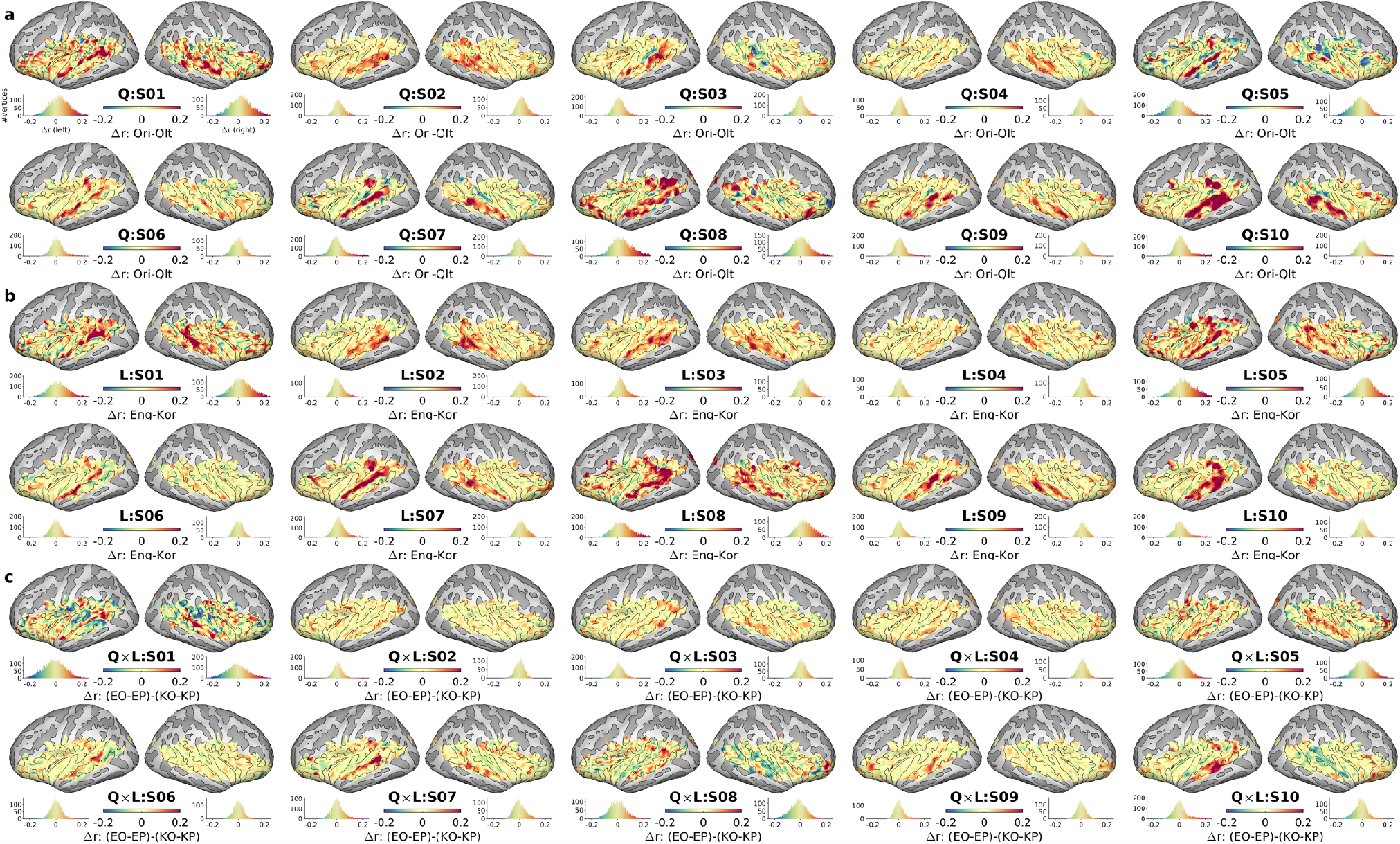
Effects of conditions on the full model prediction in individual participants. Unthresholded maps of contrasts in Pearson correlation differences are shown for (a) Quilting, (b) Language, and (c) the interaction of Quilting and Language.

**Figure S3.**
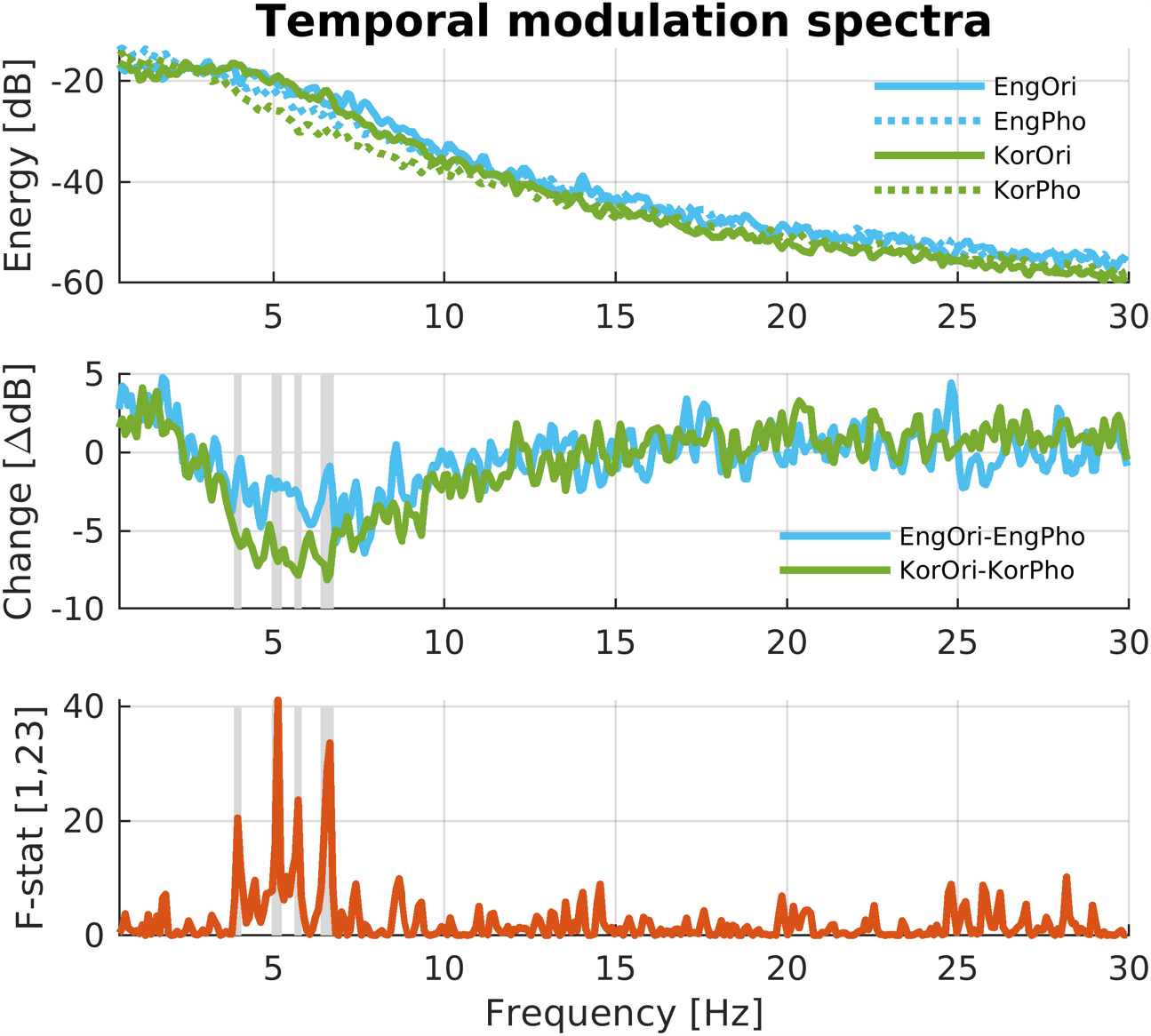
Modulation spectra of speech stimuli altered by quilting algorithm. Top: overall modulation spectra of Original (solid) and Quilts (dashed) stimuli in English (light blue) and Korean (lime green). Middle: paired differences between Original and Quilts within each language. Gray shades mark significant (FDR-P < 0.05) interaction between Quilting and Language. Bottom: F-statistics (degrees of freedom = 1, 23) from repeated measures ANOVA model for the contrast of the interaction.

**Figure S4.**
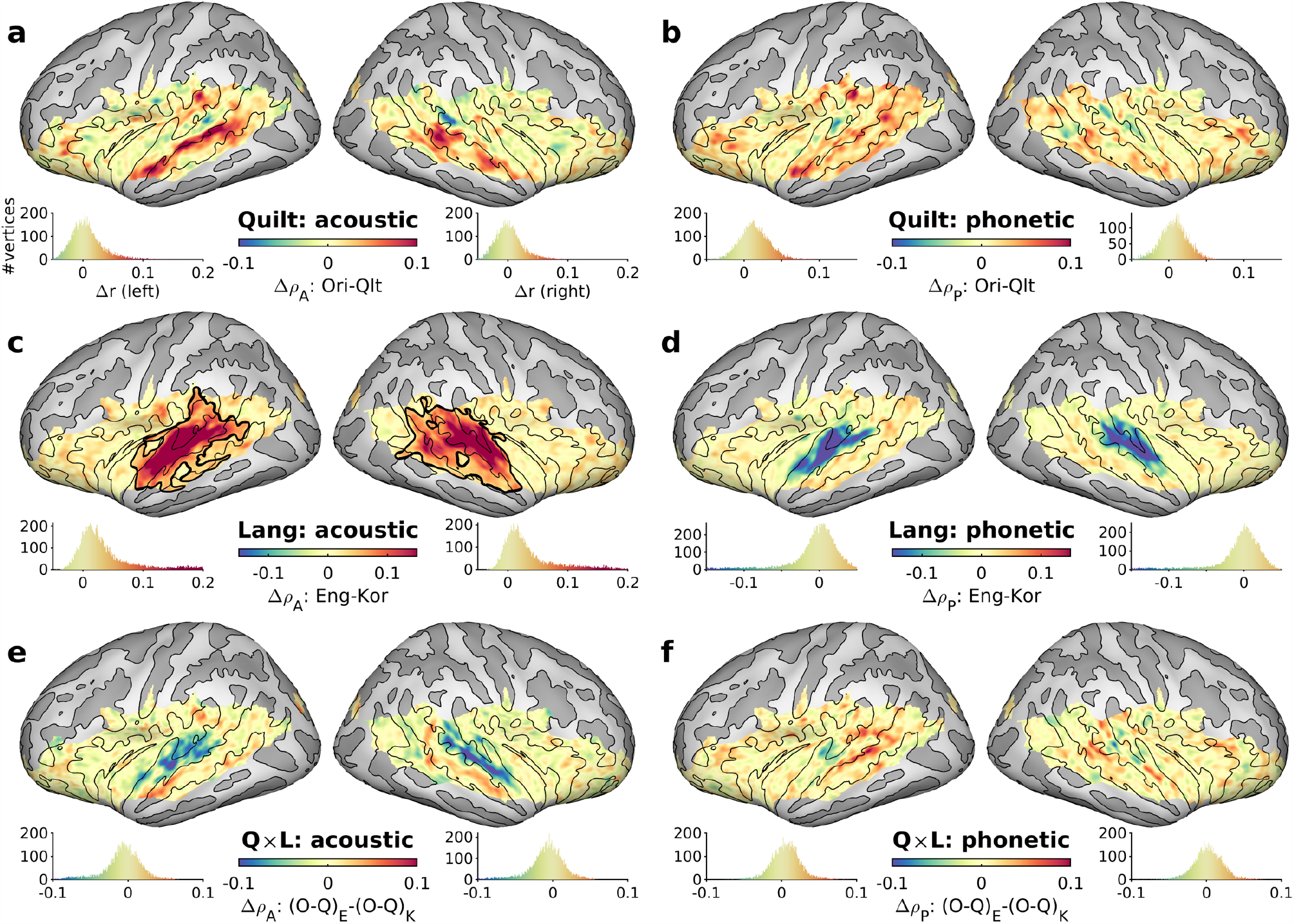
Comparisons of partial prediction accuracy between conditions: (a, b) main effect of Quilting; (c, d) main effect of Language; (e, f) interaction between Quilting and Language in partial correlation for Acoustic predictors (a, c, e) and Phonetic predictors (b, d, f). Thick black contours mark significant clusters (cluster-P ≤ 0.0025, Bonferroni adjusted alpha for the two predictor groups).

**Figure S5.**
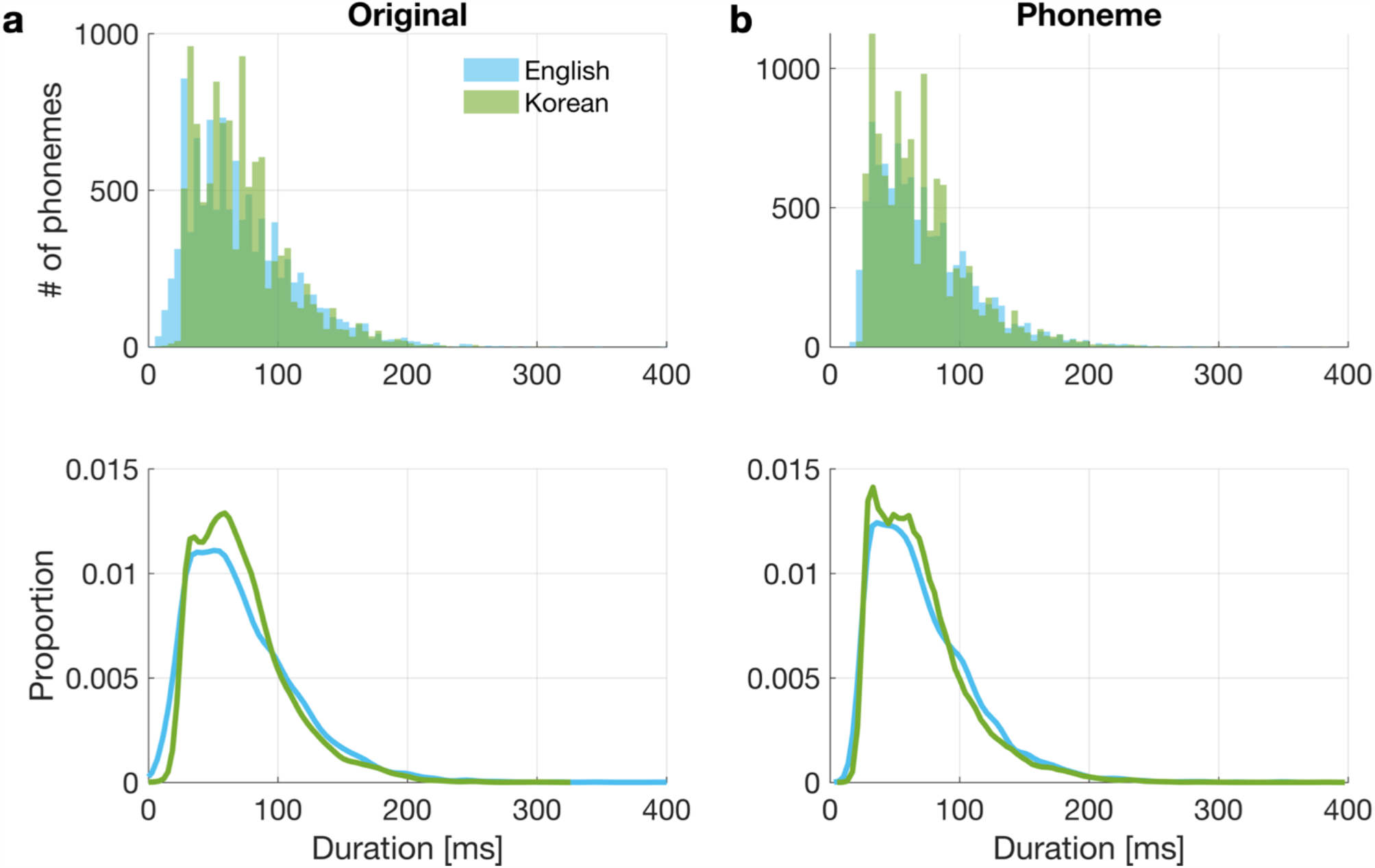
Histograms of phoneme durations. The distributions of phoneme durations in the Original natural speech (a) and the Phoneme quilts (b) are shown for English (light blue) and Korean (lime green) in histograms (top) and smoothed density functions (bottom). Non-linguistic segments (e.g., short pauses) and the last segment of each stimulus file (could have been cropped to equalize durations of stimuli) were discarded from calculation.

**Figure S6.**
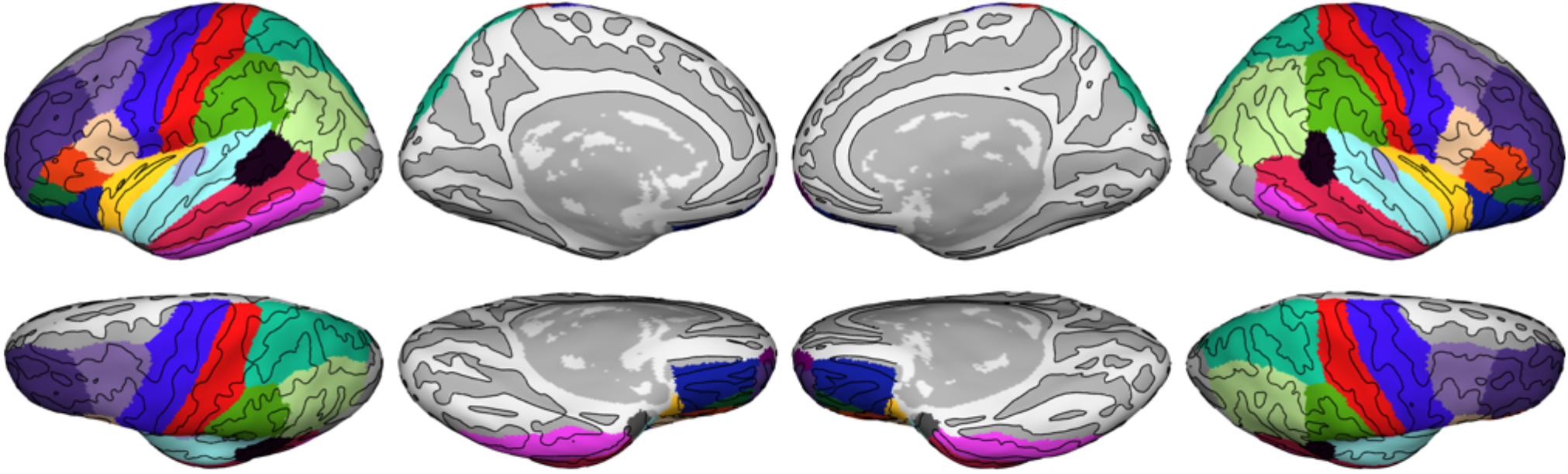
Regions of interest. From the automatic parcellation based on the Desikan-Killiany cortical atlas (Desikan et al., 2006) in FreeSurfer, the following 19 labels were included: ‘bankssts’, ‘caudalmiddlefrontal’, ‘inferiorparietal’, ‘inferiortemporal’, ‘lateralorbitofrontal’, ‘middletemporal’, ‘parsopercularis’, ‘parsorbitalis’, ‘parstriangularis’, ‘postcentral’, ‘precentral’, ‘rostralmiddlefrontal’, ‘superiorparietal’, ‘superiortemporal’, ‘supramarginal’, ‘frontalpole’, ‘temporalpole’, ‘transversetemporal’, ‘insula’.

**Figure S7.**
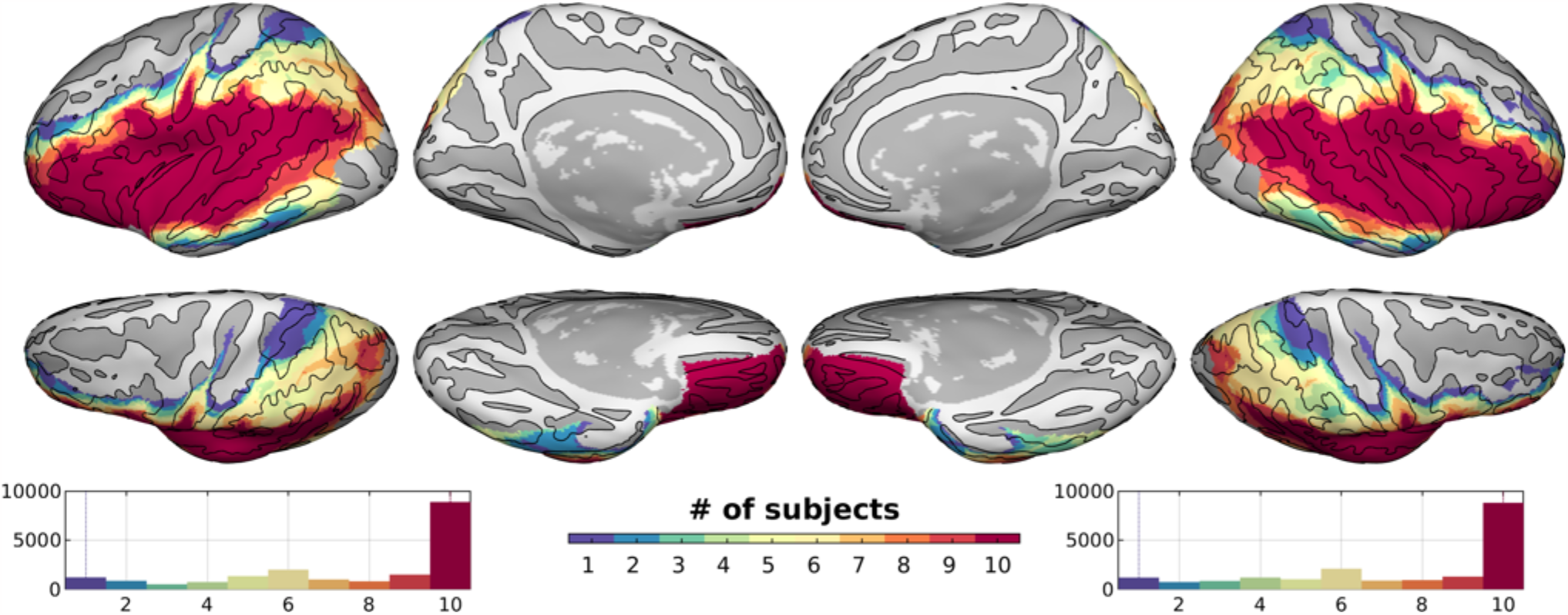
Overlap of selected vertices across participants (N = 10). The colored histograms below display the number of vertices over the number of participants.

**Figure S8.**
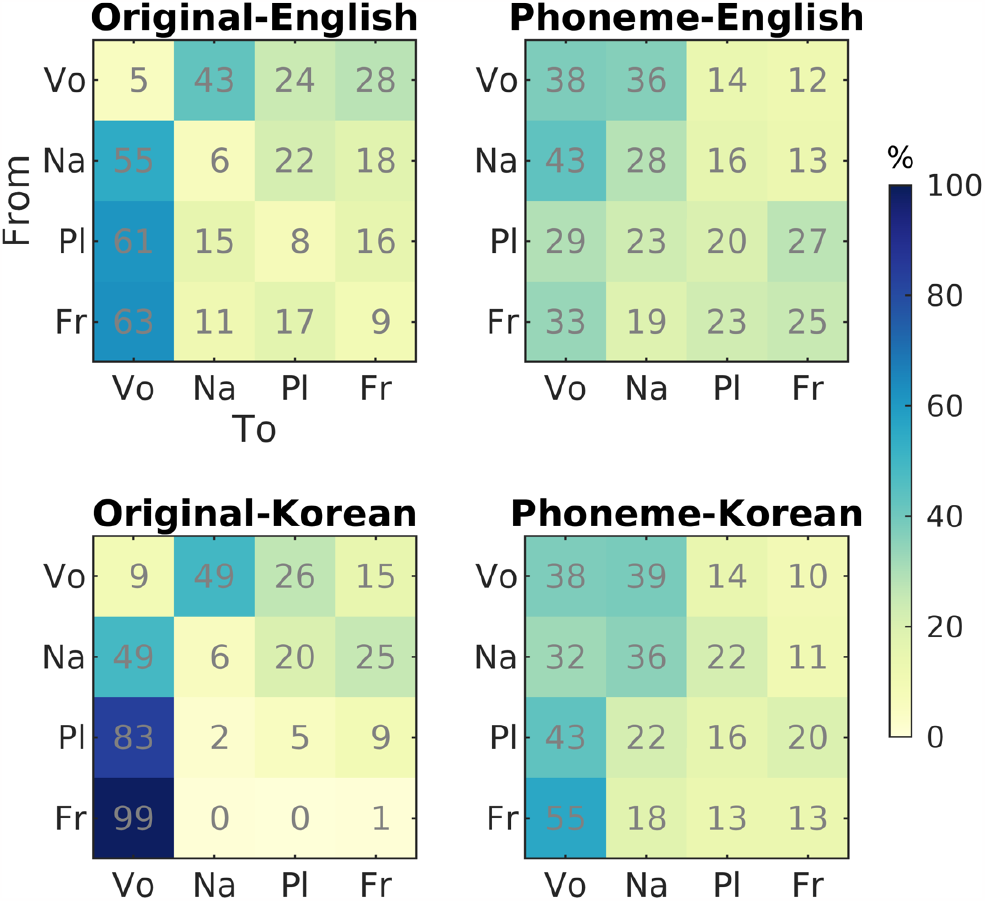
Transition probability between phoneme classes. Phoneme transitions were counted as consecutive occurrences of four phoneme classes without taking word boundaries into account, and cumulated over all stimuli for visualization. Transition probabilities are displayed from the *i*-th phoneme class (Vo, vowel; Na, nasal; Pl, plosive; Fr, fricative) in rows to the *j*-th phoneme class in columns for the four main conditions (Pr(*j*|*i*) = *T*_*i*,*j*_/Σ *T*_*i*,*j*_ where *T*_i,j_ is the number of transitions from *i* to *j*).

**Figure S9.**
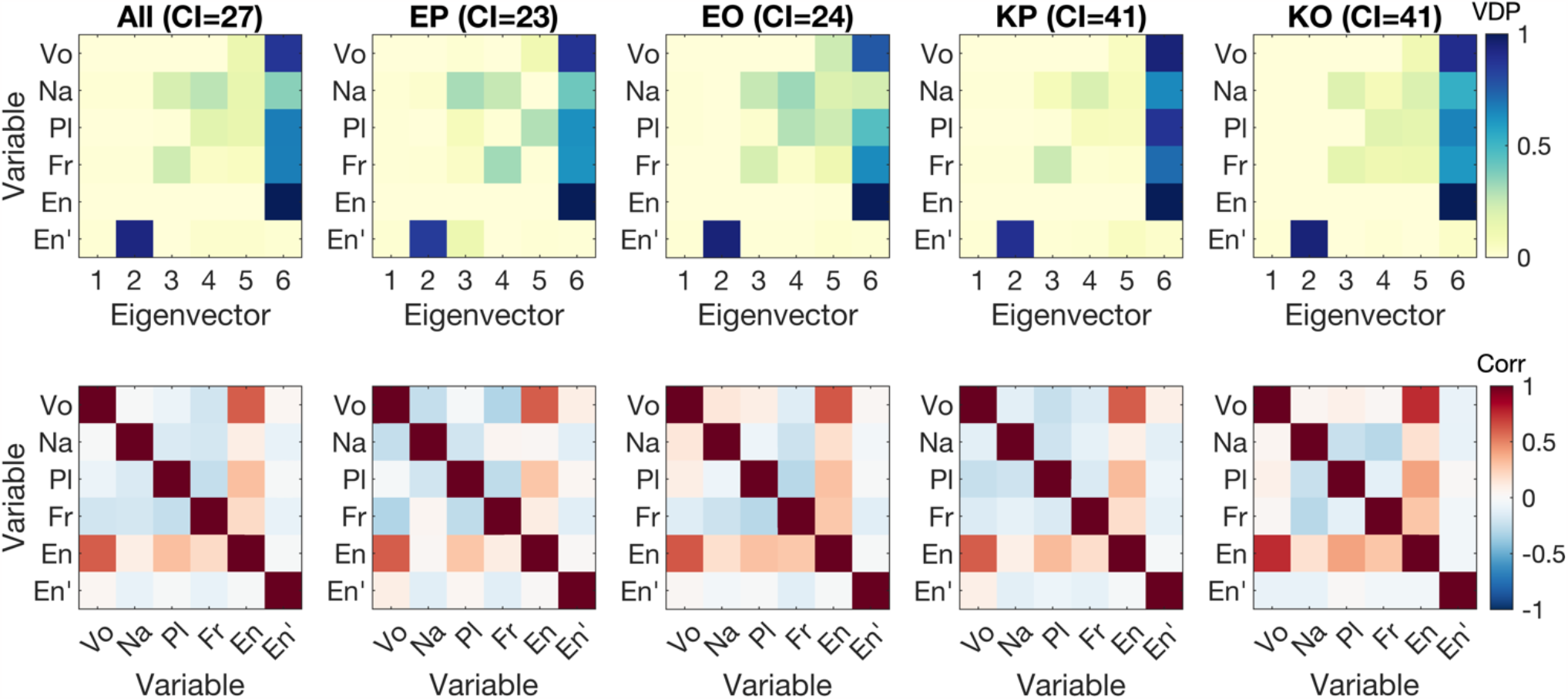
Collinearity of predictors used in the encoding analyses. Variance decomposition proportion (VDP, upper) and pair-wise Pearson correlation (corr, lower) between predictive variables (Vo, vowel; Na, nasal; Pl, plosive; Fr, fricative; En, envelope; En’, envelope derivative) are shown for all conditions together (left-most column) and for each condition (EP, English-Phoneme; EO, English-Original, KP, Korean-Phoneme; KO, Korean-Original) with conditional indices (CI).

### 7.2 Supplementary tables

**Table S1.** Individual phonemes included in analysis for each articulatory phoneme class.

|  | Vowel | Nasal/Approximant | Plosive | Fricative |
| --- | --- | --- | --- | --- |
| **English** | AA, AE, AH, AO, AW, AY, EH, ER, EY, IH, IY, OY, OW, UH, UW | L, M, N, NG, R, W, Y | B, D, G, K, P, T | CH, DH, F, JH, S, SH, TH, V, Z |
| **Korean** | A, AE, E, EO, EU, I, O, OE, U, WA, WAE, WE, WEO, WI, YA, YAE, YE, YEO, YI, YO, YU | L, M, N, NG, R | B, BB, D, DD, G, GG, K, P, T | C, H, J, JJ, S, SS |

## Footnotes

1 https://babel.ling.upenn.edu/phonetics/old_website_2015/p2fa/index.html

2 https://korean.utsc.utoronto.ca/kpa/

3 https://github.com/bbTomas/mPraat

4 http://psychtoolbox.org/

5 https://www.fil.ion.ucl.ac.uk/spm/

6 http://freesurfer.net/

7 https://fsl.fmrib.ox.ac.uk/

8 https://kendrickkay.net/GLMdenoise/

9 https://github.com/lacerbi/bads

10 http://www.fieldtriptoolbox.org/

11 https://osf.io/zgj3m/?view_only=cd4942f9ea674d79a5644796d5498e3c

## References

Acerbi, L., Ma, W.J., 2017. Practical Bayesian optimization for model fitting with Bayesian adaptive direct search. Advances in Neural Information Processing Systems 30.

Aitken, A.C., 1936. On least squares and linear combination of observations. Proceedings of the Royal Society of Edinburgh. Section B: Biology 55, 42–48.

Anderson, J.L., Morgan, J.L., White, K.S., 2003. A statistical basis for speech sound discrimination. Language and Speech 46, 155–182.

Baltzell, L.S., Srinivasan, R., Richards, V.M., 2017. The effect of prior knowledge and intelligibility on the cortical entrainment response to speech. Journal of Neurophysiology 118, 3144–3151.

Baumann, S., Joly, O., Rees, A., Petkov, C.I., Sun, L., Thiele, A., Griffiths, T.D., 2015. The topography of frequency and time representation in primate auditory cortices. Elife 4, e03256.

Behzadi, Y., Restom, K., Liau, J., Liu, T.T., 2007. A component based noise correction method (compcor) for bold and perfusion based fmri. Neuroimage 37, 90–101.

Belsley, D.A., 1991. A guide to using the collinearity diagnostics. Computer Science in Economics and Management 4, 33–50.

Blank, I.A., Fedorenko, E., 2020. No evidence for differences among language regions in their temporal receptive windows. Neuroimage 219, 116925.

Bořil, T., Skarnitzl, R., 2016. Tools rPraat and mPraat. Text, Speech, and Dialogue. TSD 2016. Lecture Notes in Computer Science. Springer International Publishing, Cham, pp. 367–374.

Breedlove, J.L., St-Yves, G., Olman, C.A., Naselaris, T., 2020. Generative feedback explains distinct brain activity codes for seen and mental images. Current Biology 30, 2211-2224.e2216.

Cheour, M., Ceponiene, R., Lehtokoski, A., Luuk, A., Allik, J., Alho, K., Näätänen, R., 1998. Development of language-specific phoneme representations in the infant brain. Nature Neuroscience 1, 351–353.

Chomsky, N., Halle, M., 1965. Some controversial questions in phonological theory. Journal of Linguistics 1, 97–138.

Cope, T.E., Sohoglu, E., Sedley, W., Patterson, K., Jones, P.S., Wiggins, J., Dawson, C., Grube, M., Carlyon, R.P., Griffiths, T.D., Davis, M.H., Rowe, J.B., 2017. Evidence for causal top-down frontal contributions to predictive processes in speech perception. Nature Communications 8, 2154.

Daube, C., Ince, R.A.A., Gross, J., 2019. Simple Acoustic Features Can Explain Phoneme-Based Predictions of Cortical Responses to Speech. Current Biology 29, 1924–1937.e1929.

Davis, M.H., Ford, M.A., Kherif, F., Johnsrude, I.S., 2011. Does semantic context benefit speech understanding through “top–down” processes? Evidence from time-resolved sparse fmri. Journal of Cognitive Neuroscience 23, 3914–3932.

Davis, M.H., Johnsrude, I.S., 2007. Hearing speech sounds: Top-down influences on the interface between audition and speech perception. Hearing Research 229, 132–147.

De Heer, W.A., Huth, A.G., Griffiths, T.L., Gallant, J.L., Theunissen, F.E., 2017. The hierarchical cortical organization of human speech processing. Journal of Neuroscience 37, 6539–6557.

Desikan, R.S., Ségonne, F., Fischl, B., Quinn, B.T., Dickerson, B.C., Blacker, D., Buckner, R.L., Dale, A.M., Maguire, R.P., Hyman, B.T., 2006. An automated labeling system for subdividing the human cerebral cortex on mri scans into gyral based regions of interest. Neuroimage 31, 968–980.

Di Liberto, G.M., O’Sullivan, J.A., Lalor, E.C., 2015. Low-frequency cortical entrainment to speech reflects phoneme-level processing. Current Biology 25, 2457–2465.

Díaz, B., Baus, C., Escera, C., Costa, A., Sebastián-Gallés, N., 2008. Brain potentials to native phoneme discrimination reveal the origin of individual differences in learning the sounds of a second language. Proceedings of the National Academy of Sciences of the United States of America 105, 16083–16088.

Ding, N., Simon, J.Z., 2013. Adaptive temporal encoding leads to a background-insensitive cortical representation of speech. The Journal of Neuroscience 33, 5728–5735.

Eckert, M.A., Teubner-Rhodes, S., Vaden, K.I., Jr., 2016. Is listening in noise worth it? The neurobiology of speech recognition in challenging listening conditions. Ear and Hearing 37, 101S–110S.

Eklund, A., Nichols, T.E., Knutsson, H., 2016. Cluster failure: Why fMRI inferences for spatial extent have inflated false-positive rates. Proceedings of the National Academy of Sciences of the United States of America, 201602413.

Friederici, A.D., 2009. Pathways to language: Fiber tracts in the human brain. Trends in Cognitive Sciences 13, 175–181.

Friederici, A.D., 2011. The brain basis of language processing: From structure to function. Physiological Reviews 91, 1357–1392.

Friederici, A.D., Pfeifer, E., Hahne, A., 1993. Event-related brain potentials during natural speech processing: Effects of semantic, morphological and syntactic violations. Cognitive Brain Research 1, 183–192.

Friederici, A.D., Wessels, J.M.I., 1993. Phonotactic knowledge of word boundaries and its use in infant speech perception. Perception and Psychophysics 54, 287–295.

Friston, K., Kiebel, S., 2009. Predictive coding under the free-energy principle. Philosophical Transactions of the Royal Society of London. Series B: Biological Sciences 364, 1211–1221.

Giraud, A.L., Kell, C., Thierfelder, C., Sterzer, P., Russ, M.O., Preibisch, C., Kleinschmidt, A., 2004. Contributions of sensory input, auditory search and verbal comprehension to cortical activity during speech processing. Cerebral Cortex 14, 247–255.

Gwilliams, L., King, J.-R., Marantz, A., Poeppel, D., 2022. Neural dynamics of phoneme sequences reveal position-invariant code for content and order. Nature Communications 13, 6606.

Hamilton, L.S., Huth, A.G., 2020. The revolution will not be controlled: Natural stimuli in speech neuroscience. Language, Cognition and Neuroscience 35, 573–582.

Hasson, U., Malach, R., Heeger, D.J., 2010. Reliability of cortical activity during natural stimulation. Trends in Cognitive Sciences 14, 40–48.

Hastie, T., Tibshirani, R., Friedman, J., 2009. The Elements of Statistical Learning: Data Mining, Inference, and Prediction. Springer Science & Business Media.

Hickok, G., Poeppel, D., 2007. The cortical organization of speech processing. Nature Reviews Neuroscience 8, 393–402.

Hoerl, A.E., Kennard, R.W., 1970. Ridge regression: Biased estimation for nonorthogonal problems. Technometrics 12, 55–67.

Holdgraf, C.R., de Heer, W., Pasley, B., Rieger, J., Crone, N., Lin, J.J., Knight, R.T., Theunissen, F.E., 2016. Rapid tuning shifts in human auditory cortex enhance speech intelligibility. Nature Communications 7, 13654.

Howard, M.F., Poeppel, D., 2010. Discrimination of speech stimuli based on neuronal response phase patterns depends on acoustics but not comprehension. Journal of Neurophysiology 104, 2500–2511.

Huth, A.G., de Heer, W.A., Griffiths, T.L., Theunissen, F.E., Gallant, J.L., 2016. Natural speech reveals the semantic maps that tile human cerebral cortex. Nature 532, 453–458.

Jusczyk, P.W., Luce, P.A., Charles-Luce, J., 1994. Infants’ sensitivity to phonotactic patterns in the native language. Journal of Memory and Language 33, 630–645.

Kaufman, S., Rosset, S., Perlich, C., Stitelman, O., 2012. Leakage in Data Mining: Formulation, Detection, and Avoidance. Acm Transactions on Knowledge Discovery from Data 6.

Kay, K., Rokem, A., Winawer, J., Dougherty, R., Wandell, B., 2013. Glmdenoise: A fast, automated technique for denoising task-based fmri data. Frontiers in Neuroscience 7.

Kay, K.N., Naselaris, T., Prenger, R.J., Gallant, J.L., 2008. Identifying natural images from human brain activity. Nature 452.

Khalighinejad, B., Cruzatto da Silva, G., Mesgarani, N., 2017. Dynamic encoding of acoustic features in neural responses to continuous speech. Journal of Neuroscience 37, 2176–2185.

Kim, S.-G., 2022. Investigating the neural encoding of musical emotion using naturalistic stimuli and computational models. figshare.

Kleinschmidt, D.F., Jaeger, T.F., 2015. Robust speech perception: Recognize the familiar, generalize to the similar, and adapt to the novel. Psychological Review 122, 148–203.

Kocagoncu, E., Clarke, A., Devereux, B.J., Tyler, L.K., 2017. Decoding the cortical dynamics of sound-meaning mapping. Journal of Neuroscience 37, 1312–1319.

Kujawa, S.G., Liberman, M.C., 2009. Adding insult to injury: Cochlear nerve degeneration after “temporary” noise-induced hearing loss. Journal of Neuroscience 29, 14077–14085.

Kumar, S., Stephan, K.E., Warren, J.D., Friston, K.J., Griffiths, T.D., 2007. Hierarchical processing of auditory objects in humans. PLoS Computational Biology 3, e100.

Kutas, M., Hillyard, S.A., 1983. Event-related brain potentials to grammatical errors and semantic anomalies. Memory and Cognition 11, 539–550.

Ladefoged, P., 2001. Vowels and Consonants : An Introduction to the Sounds of Languages. Wiley-Blackwell.

Ladefoged, P., Johnstone, K., 2015. A Course in Phonetics, Seventh edition. ed. Cengage Learning, Stamford, CT.

Leonard, M.K., Baud, M.O., Sjerps, M.J., Chang, E.F., 2016. Perceptual restoration of masked speech in human cortex. Nature Communications 7, 13619.

Lerner, Y., Honey, C.J., Silbert, L.J., Hasson, U., 2011. Topographic mapping of a hierarchy of temporal receptive windows using a narrated story. Journal of Neuroscience 31, 2906–2915.

Liberto, G.M.D., Nie, J., Yeaton, J., Khalighinejad, B., Shamma, S.A., Mesgarani, N., 2021. Neural representation of linguistic feature hierarchy reflects second-language proficiency. Neuroimage 227, 117586.

Luo, H., Poeppel, D., 2007. Phase patterns of neuronal responses reliably discriminate speech in human auditory cortex. Neuron 54, 1001–1010.

Macmillan, N.A., Kaplan, H.L., 1985. Detection theory analysis of group data: Estimating sensitivity from average hit and false-alarm rates. Psychological Bulletin 98, 185–199.

Maris, E., 2019. Enlarging the scope of randomization and permutation tests in neuroimaging and neuroscience. BioRxiv, p. 685560.

Maris, E., Oostenveld, R., 2007. Nonparametric statistical testing of EEG- and MEG-data. Journal of Neuroscience Methods 164, 177–190.

Mattys, S.L., Jusczyk, P.W., 2001. Phonotactic cues for segmentation of fluent speech by infants. Cognition 78, 91–121.

McDermott, J.H., Simoncelli, E.P., 2011. Sound texture perception via statistics of the auditory periphery: Evidence from sound synthesis. Neuron 71, 926–940.

Mesgarani, N., Chang, E.F., 2012. Selective cortical representation of attended speaker in multi-talker speech perception. Nature 485, 233–236.

Mesgarani, N., Cheung, C., Johnson, K., Chang, E.F., 2014. Phonetic feature encoding in human superior temporal gyrus. Science 343, 1006–1010.

Millman, R.E., Johnson, S.R., Prendergast, G., 2015. The Role of Phase-locking to the Temporal Envelope of Speech in Auditory Perception and Speech Intelligibility. Journal of Cognitive Neuroscience 27, 533–545.

Moerel, M., De Martino, F., Kemper, V.G., Schmitter, S., Vu, A.T., Uğurbil, K., Formisano, E., Yacoub, E., 2018. Sensitivity and specificity considerations for fmri encoding, decoding, and mapping of auditory cortex at ultra-high field. Neuroimage 164, 18–31.

Moerel, M., De Martino, F., Santoro, R., Ugurbil, K., Goebel, R., Yacoub, E., Formisano, E., 2013. Processing of natural sounds: Characterization of multipeak spectral tuning in human auditory cortex. Journal of Neuroscience 33, 11888–11898.

Moore, B.C.J., 1996. Perceptual consequences of cochlear hearing loss and their implications for the design of hearing aids. Ear and Hearing 17, 133–161.

Morosan, P., Schleicher, A., Amunts, K., Zilles, K., 2005. Multimodal architectonic mapping of human superior temporal gyrus. Anatomy and Embryology 210, 401–406.

Moulines, E., Charpentier, F., 1990. Pitch-synchronous waveform processing techniques for text-to-speech synthesis using diphones. Speech Communication 9, 453–467.

Narain, C., Scott, S.K., Wise, R.J.S., Rosen, S., Leff, A., Iversen, S.D., Matthews, P.M., 2003. Defining a left-lateralized response specific to intelligible speech using fmri. Cerebral Cortex 13, 1362–1368.

Naselaris, T., Olman, C.A., Stansbury, D.E., Ugurbil, K., Gallant, J.L., 2015. A voxel-wise encoding model for early visual areas decodes mental images of remembered scenes. Neuroimage 105, 215–228.

Norman-Haignere, S., Kanwisher, Nancy G., McDermott, Josh H., 2015. Distinct cortical pathways for music and speech revealed by hypothesis-free voxel decomposition. Neuron 88, 1281–1296.

Norman-Haignere, S.V., Long, L.K., Devinsky, O., Doyle, W., Irobunda, I., Merricks, E.M., Feldstein, N.A., McKhann, G.M., Schevon, C.A., Flinker, A., 2020. Multiscale integration organizes hierarchical computation in human auditory cortex. BioRxiv.

Nunez-Elizalde, A.O., Huth, A.G., Gallant, J.L., 2019. Voxelwise encoding models with non-spherical multivariate normal priors. Neuroimage 197, 482–492.

Obleser, J., Eisner, F., Kotz, S.A., 2008. Bilateral speech comprehension reflects differential sensitivity to spectral and temporal features. Journal of Neuroscience 28, 8116–8123.

Oganian, Y., Chang, E.F., 2019. A speech envelope landmark for syllable encoding in human superior temporal gyrus. Science Advances 5, eaay6279.

Overath, T., Lee, J.C., 2017. The neural processing of phonemes is shaped by linguistic analysis. Proceedings of the International Symposium on Auditory and Audiological Research, pp. 107–116.

Overath, T., McDermott, J.H., Zarate, J.M., Poeppel, D., 2015. The cortical analysis of speech-specific temporal structure revealed by responses to sound quilts. Nature Neuroscience 18, 903–911.

Overath, T., Paik, J.H., 2021. From acoustic to linguistic analysis of temporal speech structure: Acousto-linguistic transformation during speech perception using speech quilts. Neuroimage, 117887.

Park, H., Ince, Robin A.A., Schyns, Philippe G., Thut, G., Gross, J., 2015. Frontal top-down signals increase coupling of auditory low-frequency oscillations to continuous speech in human listeners. Current Biology 25, 1649–1653.

Poeppel, D., Idsardi, W.J., Wassenhove, V.v., 2008. Speech perception at the interface of neurobiology and linguistics. Philosophical Transactions of the Royal Society B: Biological Sciences 363, 1071–1086.

Rao, R.P.N., Ballard, D.H., 1999. Predictive coding in the visual cortex: A functional interpretation of some extra-classical receptive-field effects. Nature Neuroscience 2, 79–87.

Rauschecker, J.P., Scott, S.K., 2009. Maps and streams in the auditory cortex: Nonhuman primates illuminate human speech processing. Nature Neuroscience 12, 718–724.

Rauschecker, J.P., Tian, B., 2004. Processing of band-passed noise in the lateral auditory belt cortex of the rhesus monkey. Journal of Neurophysiology 91, 2578–2589.

Ringach, D.L., Sapiro, G., Shapley, R., 1997. A subspace reverse-correlation technique for the study of visual neurons. Vision Research 37, 2455–2464.

Ruggles, D., Bharadwaj, H., Shinn-Cunningham, B.G., 2011. Normal hearing is not enough to guarantee robust encoding of suprathreshold features important in everyday communication. Proceedings of the National Academy of Sciences of the United States of America 108, 15516–15521.

Rutten, S., Santoro, R., Hervais-Adelman, A., Formisano, E., Golestani, N., 2019. Cortical encoding of speech enhances task-relevant acoustic information. Nature Human Behaviour 3, 974–987.

Saenz, M., Langers, D.R.M., 2014. Tonotopic mapping of human auditory cortex. Hearing Research 307, 42–52.

Saffran, J.R., Newport, E.L., Aslin, R.N., 1996. Word segmentation: The role of distributional cues. Journal of Memory and Language 35, 606–621.

Samuel, A.G., 1981. Phonemic restoration: Insights from a new methodology. Journal of Experimental Psychology: General 110, 474–494.

Samuel, A.G., 1987. Lexical uniqueness effects on phonemic restoration. Journal of Memory and Language 26, 36–56.

Santoro, R., Moerel, M., De Martino, F., Goebel, R., Ugurbil, K., Yacoub, E., Formisano, E., 2014. Encoding of natural sounds at multiple spectral and temporal resolutions in the human auditory cortex. PLoS Computational Biology 10.

Santoro, R., Moerel, M., De Martino, F., Valente, G., Ugurbil, K., Yacoub, E., Formisano, E., 2017. Reconstructing the spectrotemporal modulations of real-life sounds from fmri response patterns. Proceedings of the National Academy of Sciences of the United States of America 114, 4799–4804.

Scott, S.K., Blank, C.C., Rosen, S., Wise, R.J.S., 2000. Identification of a pathway for intelligible speech in the left temporal lobe. Brain 123, 2400–2406.

Shannon, R.V., Zeng, F.-G., Kamath, V., Wygonski, J., Ekelid, M., 1995. Speech recognition with primarily temporal cues. Science 270, 303–304.

Shin, J., 2015. Vowels and consonants. In: Brown, L., Yeon, J. (Eds.), The handbook of Korean linguistics. Wiley-Blackwell, UK, pp. 3–21.

Shinn-Cunningham, B.G., Best, V., 2008. Selective attention in normal and impaired hearing. Trends in Amplification 12, 283–299.

Sohn, H.-M., 2001. The Korean Language. Cambridge University Press, NY.

Sohoglu, E., Peelle, J.E., Carlyon, R.P., Davis, M.H., 2012. Predictive top-down integration of prior knowledge during speech perception. Journal of Neuroscience 32, 8443–8453.

Stevens, K.N., 2000. Acoustic Phonetics. MIT press.

Theunissen, F., Miller, J., 1995. Temporal encoding in nervous systems: A rigorous definition. Journal of Computational Neuroscience 2, 149–162.

van de Wiel, M.A., van Nee, M.M., Rauschenberger, A., 2021. Fast Cross-validation for Multi-penalty High-dimensional Ridge Regression. Journal of Computational and Graphical Statistics 30, 835–847.

Vanthornhout, J., Decruy, L., Wouters, J., Simon, J.Z., Francart, T., 2018. Speech Intelligibility Predicted from Neural Entrainment of the Speech Envelope. Journal of the Association for Research in Otolaryngology 19, 181–191.

Varoquaux, G., Raamana, P.R., Engemann, D.A., Hoyos-Idrobo, A., Schwartz, Y., Thirion, B., 2017. Assessing and tuning brain decoders: Cross-validation, caveats, and guidelines. Neuroimage 145, 166–179.

Verschueren, E., Vanthornhout, J., Francart, T., 2021. The effect of stimulus intensity on neural envelope tracking. Hearing Research 403, 108175.

Warren, J.D., Jennings, A.R., Griffiths, T.D., 2005. Analysis of the spectral envelope of sounds by the human brain. Neuroimage 24, 1052–1057.

Warren, R.M., 1970. Perceptual restoration of missing speech sounds. Science 167, 392–393.

Wild, C.J., Davis, M.H., Johnsrude, I.S., 2012. Human auditory cortex is sensitive to the perceived clarity of speech. Neuroimage 60, 1490–1502.

Wu, M.C.-K., David, S.V., Gallant, J.L., 2006. Complete functional characterization of sensory neurons by system identification. Annual Review of Neuroscience 29, 477–505.

Yi, H.G., Leonard, M.K., Chang, E.F., 2019. The encoding of speech sounds in the superior temporal gyrus. Neuron 102, 1096–1110.

Yoon, T.-J., Kang, Y., 2013. The Korean Phonetic Aligner Program Suite.

Yuan, J., Liberman, M., 2008. Speaker identification on the scotus corpus. Journal of the Acoustical Society of America 123, 3878.

